# Multi-Omic Analyses of Dietary Fatty Acid-Microbe-Host Interactions Reveal Metaorganismal Lipid Metabolic Crosstalk Impacting Cardiometabolic Disease

**DOI:** 10.64898/2026.02.16.705588

**Authors:** Nour Mouannes, Amy C. Burrows, Anthony J. Horak, Venkateshwari Varadharajan, Sumita Dutta, Kala Mahen, Rakhee Banerjee, William J. Massey, Xiayan Ye, Marko Mrdjen, Amanda L. Brown, Olumuyiwa Awoniwi, Kohey Kitao, Adarsh Sandhu, Chiaki Tomimoto, Kowa Tsuji, Yasunori Yonejima, Isaac Ampong, Renliang Zhang, Yunguang Qiu, Belinda Willard, Adeline M. Hajjar, Mohammed Dwidar, Naseer Sangwan, Mary E. Walker, Matthew Spite, Feixiong Cheng, J. Mark Brown

## Abstract

Following a meal, our gut microbiome and human cells collaborate via metaorganismal metabolic circuits to produce diverse nutrient metabolites that systemically circulate to influence health and disease. Although there are now several examples of bacterial fiber-, amino acid-, and micronutrient-derived metabolites impacting cardiometabolic disease, very little is known in regards to how diet-microbe-host interactions impact lipid homeostasis. Here we address this by defining dietary fatty acid substrate availability in germ-free versus conventionally-raised mice coupled to deep multi-omic metabolic phenotyping. Our data demonstrate that the effects of dietary saturated (SFA), monounsaturated (MUFA), and polyunsaturated fatty acids (PUFA) on the host lipidome, transcriptome, proteome and metabolome are uniquely impacted by resident microbiota. Also, the hepatic levels of both pro-inflammatory and pro-resolving lipid mediators are strongly influenced by dietary fatty acid-microbe interactions. This study presents a unique resource to the nutrition and metabolism research community to advance our understanding of metaorganismal lipid metabolism.

GRAPHICAL ABSTRACT

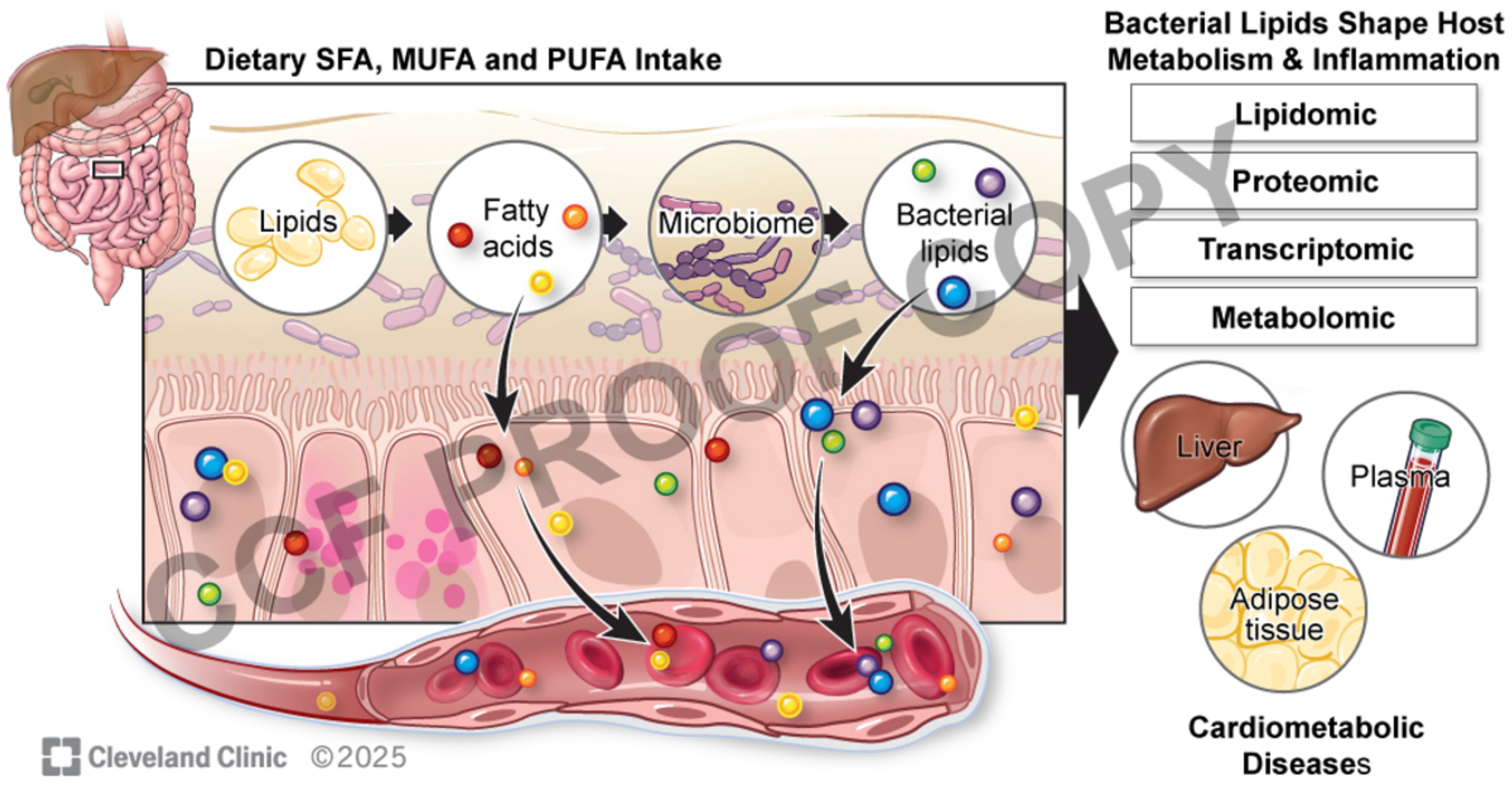

## INTRODUCTION

A common hallmark of cardiometabolic diseases (CMD) such as obesity, type 2 diabetes mellitus (T2DM), and atherosclerotic cardiovascular disease (ASCVD) is low-grade chronic inflammation^1–3^. Emerging evidence suggests that CMD-associated inflammation is strongly influenced by diet-microbe-host-interactions^4–10^. In fact, nearly all CMD-related comorbidities including obesity, insulin resistance, and atherosclerosis can be transferred into naïve germ-free (GF) mice by fecal microbial transplantation^4–10^. Clearly, the gut microbiome plays a key role in diet-driven pathways promoting these common diseases of unresolved inflammation. However, the mechanisms linking diet-microbe-host interactions to CMD remain poorly understood. One prevailing hypothesis is that poor nutrition and associated obesity dually promote gut “dysbiosis” and increase intestinal permeability to allow for the translocation of microbe-associated molecular patterns (MAMPs) such as lipopolysaccharide (LPS) into the circulation^11–13^. MAMPs can then engage with host pattern recognition receptors such as members of the toll-like receptors (TLR) family expressed by resident macrophages of many tissues to promote metabolic inflammation^11–13^. Although there is some support for this “metabolic endotoxemia” hypothesis, the role for circulating gut microbe-derived metabolites as key regulators of CMD is also being increasingly appreciated. The alternative gut microbial endocrine organ hypothesis^14–16^ posits that gut bacteria can produce meal-related metabolites that engage with host receptors to impact CMD. This metaorganismal endocrinology hypothesis^14–16^ differs from the classic pattern recognition paradigm, and instead focuses on how both gut bacteria and our human cells coordinate nutrient metabolism to impact disease. Here, we have specifically interrogated how the metaorganismal metabolism of dietary fatty acids impact CMD in the host, given the central role that fatty acids play in the production of signaling lipid mediators that either promote inflammation or facilitate the resolution of inflammation^17–20^. Beyond their role as substrates for the dynamic generation of pro-inflammatory and pro-resolving lipid mediators^17–20^, dietary fatty acids also serve as key cellular energy sources, building blocks for membrane phospholipids and stored triglycerides, and signaling molecules for both our human cells and bacteria living within us^21^. Although we have a wealth of knowledge regarding lipid metabolism in mammalian cells, there is very little known regarding commensal microbial fatty acid metabolism, and even less known about microbe-host integration in metaorganismal lipid metabolic circuits.

It is important to remember that one of the most striking predictors of CMD-related mortality from many independent large epidemiological studies is the type of dietary fats being consumed (*i.e.* high dietary saturated fat and cholesterol are strongly correlated with adverse outcomes)^22–28^. There is overwhelming evidence that replacing diets high in saturated fatty acids (SFA) with diets containing high ω-6 or ω-3 PUFAs can slow progression of CMD in rodents and humans^29–35^. Most recently, randomized controlled trials have shown that pharmaceutical grade ω-3 PUFA supplementation can improve CMD outcomes^32–35^. The REDUCE-IT (Reduction of Cardiovascular Events with Icosapent Ethyl-Intervention Trial) clinical trial showed that icosapent ethyl (a stable eicosapentaenoic acid derivative) significantly reduced major adverse cardiovascular events^32–35^. Icosapent ethyl has also shown benefit in related CMD phenotypes including obesity, insulin resistance, and fatty liver disease^32–35^. However, not all studies have shown clear benefit from icosapent ethyl^34,35^, and this variable response has caused some skepticism in the community whether ω-3 PUFAs are a viable treatment option for CMD. Here, we hypothesized that the ability of select ω-3 and ω-6 PUFAs to improve CMD relies in part on alterations in the gut microbiome community and related production of PUFA-derived microbial metabolites that impact host lipid metabolism, inflammatory responses, and CMD. To rigorously test this hypothesis we fed either GF or conventionally housed mice well-defined sterile high fat diets containing either plant- or animal-derived SFA, monounsaturated fatty acids (MUFA), or ω-3 and ω-6 PUFAs and performed a multi-omic investigation of CMD-relevant phenotypes.

Collectively, our data demonstrate that gut microbe-driven metabolism of dietary PUFAs is required for some of the remodeling of membrane phospholipids, oxylipin production, and suppression of CMD in the host. This study provides a multi-omic resource allowing for discovery of unique dietary fatty acid-microbe-host interactions that impact metabolic physiology relevant to CMD and many other fields related to unresolved inflammation.

## RESULTS

### The Ability of Dietary Fatty Acids To Impact Metabolic Physiology is Shaped By Resident Microbiota

The ability of dietary SFA, MUFA and PUFA supplementation to differentially impact CMD is well established^22–35^. However, an important gap exists as to how dietary SFA, MUFA, and PUFA substrates interact with the gut microbiome to influence systemic lipid metabolism and metabolic physiology within the broader metaorganism. Unfortunately, most animal studies rely on standard rodent “chow” formulations which are grain-based and very high in fiber, low fat, and the composition of the macronutrients including fatty acids are poorly defined. In fact, chow-based formulations vary substantially by batch which limit rigor and reproducibility in affecting metabolic phenotypes in mice^36^. To overcome these key limitations, and to study obesity-related CMD phenotypes, we synthesized chemically defined obesogenic diets with well-defined SFA, MUFA, and PUFA levels (Figure 1A,1B; Supplemental Table 1). Given we wanted to study the impact of dietary fatty acids under GF settings, we also ensured all experimental diets were sterile (i.e. devoid of any live bacteria) by double irradiation, sterile vacuum packaging, and careful sterile introduction to all mouse cages. The control obesogenic diet (20% protein, 40% carbohydrate, and 40% fat) contained added fructose, high cholesterol, and high fat with a base of palm oil, which is mostly enriched in SFA including palmitate (16:0) and stearate (18:0) (PALM, Figure 1A,1B; Supplemental Table 1). To provide high MUFA substrate, we created a calorically balanced diet that contained high oleic acid (18:1, ω-9) safflower oil in the place of the palm oil base in the control diet (HOS, Figure 1A,1B; Supplemental Table 1). Next, to provide both plant and animal sources of defined ω-6 PUFA substrates we replaced the palm oil base with either high linoleic acid (LA, 18:2, ω-6) safflower oil (HLS, Figure 1A,1B; Supplemental Table 1) or γ-linolenic acid (GLA, 18:3, ω-6)-enriched borage oil (BOR, Figure 1A,1B; Supplemental Table 1).

**Figure 1.**
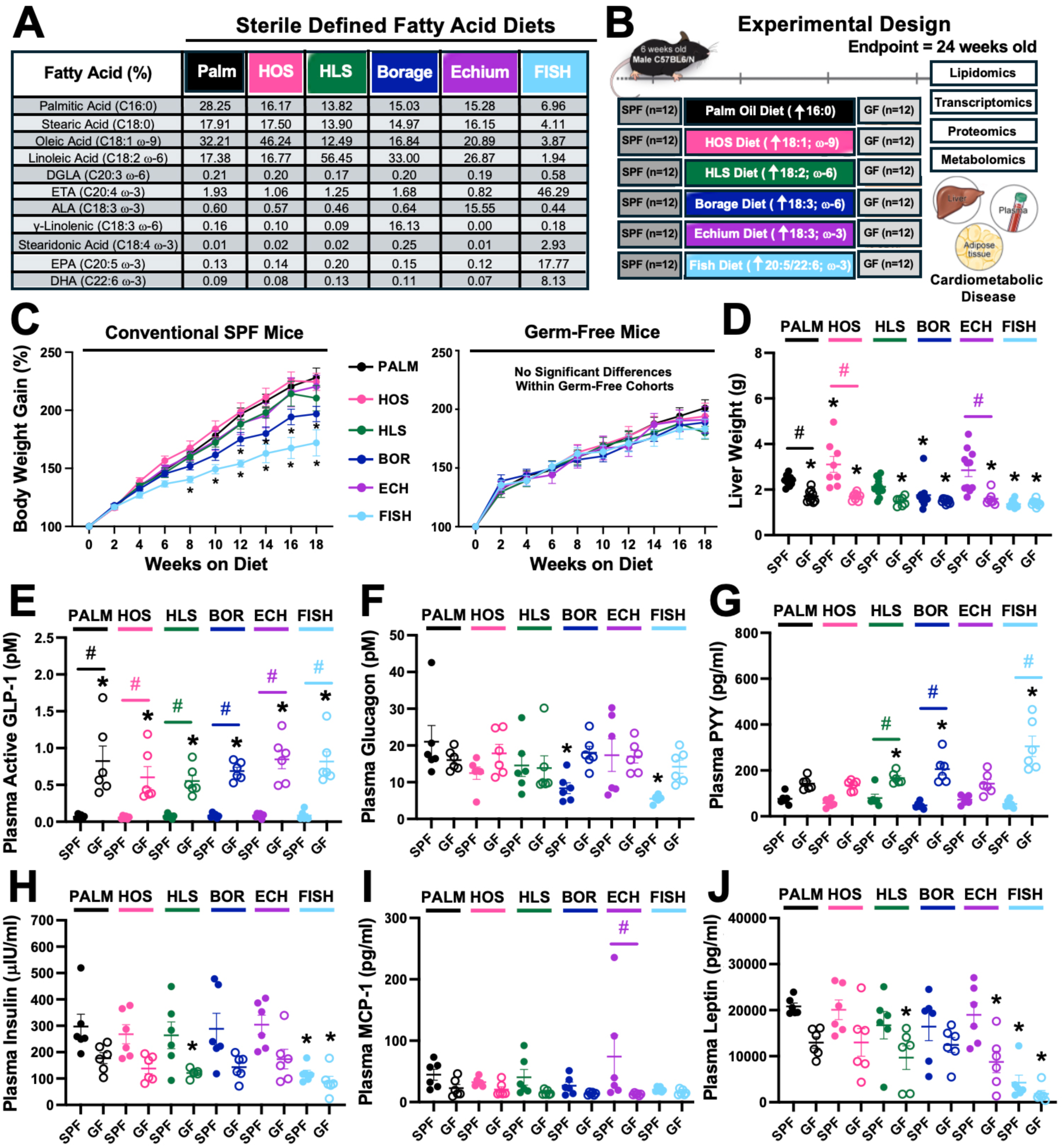
The Ability of Dietary Fatty Acids to Alter Metabolic Physiology is Shaped by Resident Microbiota. **(A)** The fatty acid composition (expressed as a % of the total fatty acid) of all sterile experimental diets was quantified using liquid chromatography tandem mass spectrometry (LC-MS/MS). **(B)** Overview of the experimental design to feed sterile diets with well-defined fatty acid composition to either specific pathogen-free (SPF) conventionally housed mice or mice maintained under germ-free conditions (GF). Either SPF or GF cohorts of male C57BL/6N mice were maintained on sterile fatty acid-defined diets containing saturated (Palm), high oleic safflower (HOS), high linoleic safflower (HLS), borage (BOR), echium (ECH) or fish oils for 18 weeks. Resulting samples including plasma, intestine, liver, and adipose tissue were then subjected to extensive lipidomic, transcriptomic, proteomic, and metabolomic analyses to understand diet-microbe-host interactions in lipid metabolism. **(C)** Body weight gain (%) comparison between conventional SPF and GF mice. **(D)** Liver weight (grams) following necropsy. **(E-F)** Plasma levels metabolic hormones and chemokines including **(E)** glucagon-like peptide 1 (GLP-1), **(F)** glucagon, **(G)** peptide YY (PYY), **(H)** insulin, **(I)** monocyte chemoattractant protein 1 (MCP-1), and **(J)** leptin were quantified using multiplex immunoassays. Data represent the mean ± S.E.M. from n=6-12 per group, and statistically significant differences (*p*<0.05) were detected using ANOVA with post-hoc Tukey-Kramer HSD for all pairs comparisons. * = significantly different than the SPF palm oil-fed control group; # = significantly different when comparing SPF and GF groups within each dietary condition.

Also, to provide plant and animal sources of defined ω-3 PUFA substrates we created matched diets with either α-linolenic acid (ALA, 18:3, ω-3)-enriched echium oil (ECH, Figure 1A,1B; Supplemental Table 1), or fish oil which has high levels of both eicosapentaenoic acid (EPA, 20:5, ω-3) and docosahexaenoic acid (DHA, 22:6, ω-3) (FISH, Figure 1A,1B; Supplemental Table 1). These sterile chemically-defined diets were then fed to either conventionally housed specific pathogen-free (SPF) mice with an intact microbiome or mice maintained under germ-free (GF) conditions for a period up to 18 weeks (Figure 1B). Leveraging this rigorous experimental approach, we were able to deeply phenotype metabolic physiology and systemic lipid metabolism to uncover many unexpected dietary fatty acid-microbe-host interactions.

One of the first diet-microbe-host interactions we noticed was a clear difference in the ability of different dietary fats to impact obesity-related phenotypes (Figure 1). For instance, in conventional SPF mice both borage and fish oil groups gained less body weight compared to the palm oil control group (Figure 1C). However, the ability of dietary fatty acid composition to alter body weight gain was absent in GF mice (Figure 1C). Also, conventionally-raised SPF mice had increased liver weight with HOS feeding, yet reduced liver weight with either borage or fish oil feeding, compared to the palm oil control (Figure 1D). In contrast, all GF mice had the same liver weight regardless of which dietary substrate was provided (Figure 1D). We next quantified the circulating levels of several key metabolic hormones and cytokines/chemokines, and identified clear diet-microbe-host interactions (Figure 1E-1J). In agreement with previous reports^37^, GF mice have much higher levels of active glucagon-like peptide 1 (GLP-1), but this effect was very similar across all diet groups (Figure 1E). In contrast, only in conventional SPF cohorts, both borage and fish oil groups had reduced glucagon levels compared to the palm oil controls (Figure 1F). Furthermore, the appetite-regulating hormone peptide YY (PYY) was found to be elevated in all GF cohorts compared to SPF controls (Figure 1G). However, the elevated levels of PYY seen in GF mice was significantly enhanced in mice consuming fish oil when compared to all other groups (Figure 1G). Although plasma insulin levels were not significantly altered between SPF and GF cohorts within each diet group, GF mice fed either HLS or fish had reduced insulin levels compared to all other groups (Figure 1H). The chemokine monocyte chemoattractant 1 (MCP-1) was not strongly altered by dietary fatty acid availability, but was significantly reduced in GF mice compared to SPF mice only when fed echium oil (Figure 1I). Also, circulating leptin levels were lowest in mice consuming fish oil, but also trended to be lower in all GF cohorts compared to SPF mice in a diet-specific manner (Figure 1J).

We next examined the ability of dietary fatty acid composition to alter host gene expression in the colon, liver, adipose tissue in SPF and GF mice (Figures S1 and S2A). First, we performed bulk RNA-seq in colon (i.e. where microbiota reside in high abundance) and then performed pathway enrichment on this data using the Reactome database as the reference database, and discovered that there were clear diet-microbe-host interactions in host gene expression (Figure S2A). As expected, in the colon the major pathways affected differentially in SPF and GF mice when fed defined fatty acid substrates were fatty acid and phospholipid metabolism pathways (Figure S2A). In addition, other pathways showing clear diet-microbe-host interactions in the colon included toll-like receptor (TLR) signaling, translational control, and several diverse receptor signaling pathways including MET proto-oncogene (MET), epidermal growth factor receptor (EGFR), transforming growth factor β (TGF-β), and Notch (Figure S2A). We next evaluated the expression of host lipogenic genes in the colon, liver, and gonadal white adipose tissue (gWAT) and found tissue-specific alterations (Figure S1). In the liver, lipogenic genes including sterol regulatory element-binding protein 1c (*Srebf1*), acetyl-CoA carboxylase 1 α (*Acaca*), and fatty acid synthase (*Fasn*) were generally elevated in GF mice compared to SPF mice in the control palm oil-fed group, but reciprocally reduced in GF versus SPF in mice consuming w-3 substrates from echium or fish oil (Figure S1A-S1C). In contrast, in gWAT the expression of *Srebf1*, *Acaca*, and *Fasn* were not significantly different comparing GF and SPF cohorts, but were altered in a dietary fatty acid substrate-dependent manner (Figure S1E-S1G). However, in the colon the expression of lipogenic genes was not dramatically altered by either the presence of microbiota (i.e. comparing GF to SPF) or by diet (Figure S1I-S1K). Also, in general the expression of the macrophage marker gene CD68 (*Cd68*) did not show clear differential abundance across groups, but it is interesting to note that conventional mice fed 18:2;ω-6-enriched HLS has elevated CD68 expression in the liver and gWAT only in conventional SPF mice, but not in GF mice, when compared to the palm oil control group (Figure S1D-S1L). Collectively, our findings suggest that some, but not all, of the effects of dietary fatty acid composition on body weight, liver weight, circulating metabolic hormones, and host gene expression is shaped by resident microbiota (Figures 1, S1, and S2A).

### Dietary Fatty Acid Composition Uniquely Alters the Gut Microbiome

We next examined diet-driven alterations in the cecal microbiome to more broadly understand how dietary fatty acid composition impact the gut microbiome in conventional mice (Figure 2). First, we confirmed that our irradiated experimental diets were sterile (i.e. devoid of living bacteria) as it is well known that casein in special diets yields a positive signal from abundant *Lactococcus*^38^. To confirm sterility, we fed each diet to control germ-free mice for 1 week and found a strong 16S rRNA PCR signal from fecal DNA (Figure 2A). However, cultures were negative, and 16S rRNA PCR signals vanished when the mice were switched back to autoclaved chow for either 2 or 4 weeks (Figure 2A). When we next examined the cecal microbiome of mice fed our sterile defined fatty acid diets for 18 weeks (Figure 2B-2E). It is important to acknowledge that all groups had detectable 16S reads at endpoint (Figure 2A,2B), given it is impossible to synthesize diets without any resident bacteria (i.e. the oils and other raw material incorporated into the diets have resident bacteria). However, the GF cohorts showed marked reduction in *Phylum*-level alpha diversity compared to SPF groups (Figure 2B). When comparing all groups, *Phylum*-level analysis revealed that *Firmicutes* (68.8%–75.3%) and *Bacteroidota* (16.5%– 22.2%) were the dominant taxa across all experimental groups (Figure 2B). While the SPF palm, HOS, HLS, borage, and echium groups maintained largely comparable phylum compositions, the fish oil group exhibited notable shifts. Specifically, a substantial decrease in *Deferribacterota* (2.93%) compared to the plant oil groups (3.65%–6.59%). Concurrently, *Actinobacteriota* (0.0672%), and *Desulfobacterota* (1.56%) were generally elevated in the fish group, further differentiating its community structure and underpinning the observed reduction in alpha diversity (Figure. 2B).

**Figure 2.**
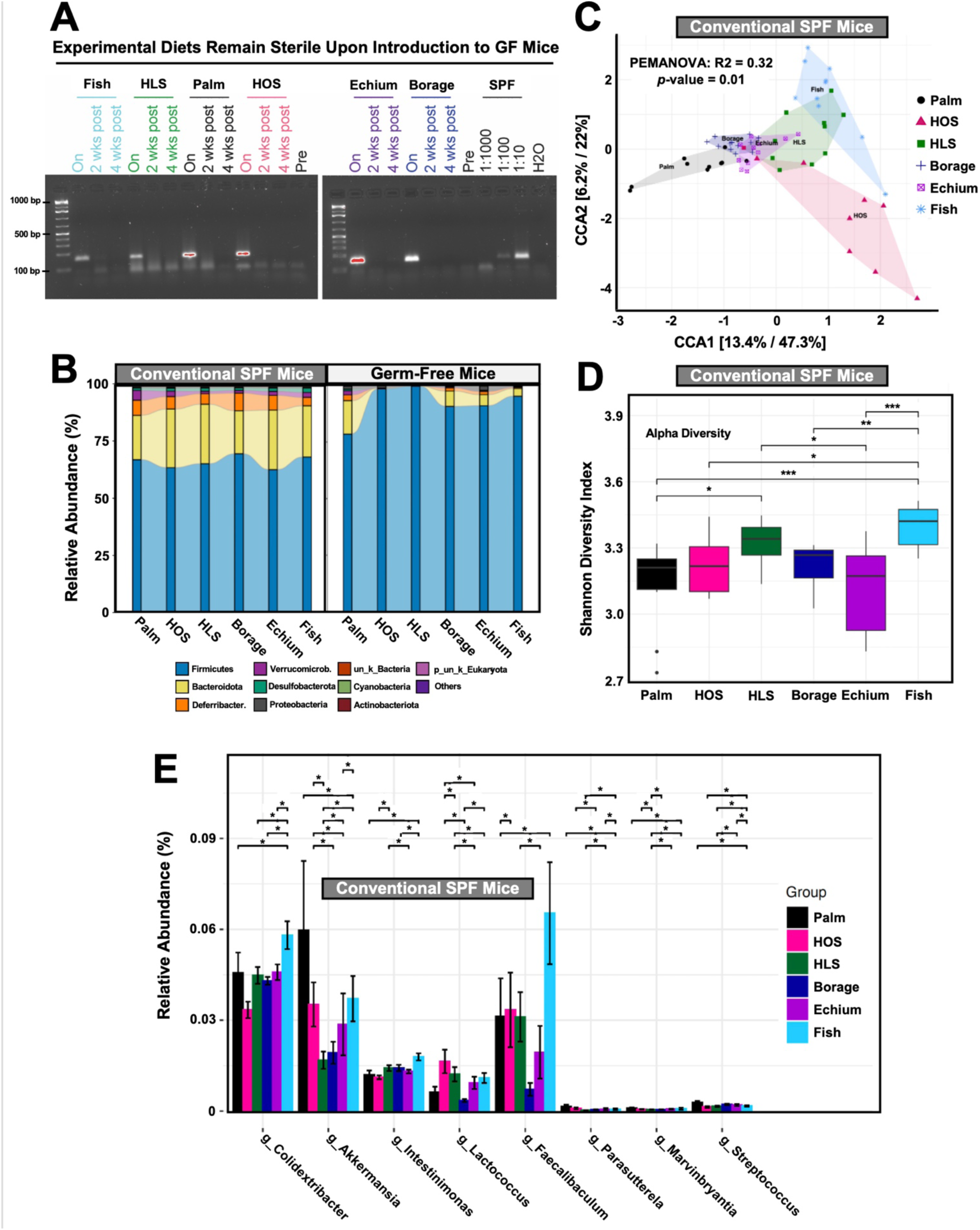
Dietary Fatty Acid Composition Alters the Gut Microbiome. Starting at 6 weeks of age, either conventionally-raised specific pathogen-free (SPF) or germ-free (GF) cohorts of male C57BL/6N mice were maintained on sterile fatty acid-defined diets containing saturated (Palm), high oleic safflower (HOS), high linoleic safflower (HLS), borage (BOR), echium (ECH) or fish oils for up to 18 weeks. **(A)** To ensure sterility of the experimental diets, a subset of GF mice were provided with each diet for 1 week and then switched back to autoclaved chow for 4 weeks. Fecal samples before the special diet (Pre), after 1 week of feeding the special diet (On), and then at 2 and 4 weeks post switch back to autoclaved chow. 16S rRNA PCR products from DNA extracted from the fecal samples were separated on a 2% agarose gel. The expected size product of 147 bp is indicated by the arrow. Fecal DNA extracted from an SPF mouse was used as a positive control. **(B)** Cecal microbiome composition analyses were done via sequencing the V4 region of the 16S rRNA the relative abundance at the *phylum* taxonomic level are shown for both conventional SPF and GF housed groups. **(C)** Canonical correspondence analysis (CCA) based beta diversity analyses in conventional SPF mice show distinct microbiome compositions in diet groups. Statistical significance and beta dispersion were estimated using PERMANOVA. **(D)** Alpha diversity (Shannon diversity index) analyses in conventional SPF mice. Each box represents the interquartile range (IQR) for a diet group, with the horizontal line indicating the median. Whiskers extend to the lowest and highest data points within 1.5 times the IQR, while individual data points are overlaid as jittered dots. The y-axis shows Shannon diversity index values, with higher values indicating greater diversity. Statistical significance between groups is denoted by connecting lines with asterisks (* p < 0.05, ** p < 0.01, *** p < 0.001) based on pairwise Wilcoxon tests with Benjamini-Hochberg correction, showing only comparisons with p < 0.1. **(E)** Significantly altered cecal microbial genera across groups using the metagenomeSeq method. The x-axis represents the log2 fold change in genus abundance, while the y-axis lists the top 10 differentially abundant genera. Statistically significant differences (p < 0.05) are indicated by asterisks.

To assess the impact of dietary fatty acid composition in SPF cohorts on overall microbial community structure, a Permutational Multivariate Analysis of Variance (PERMANOVA) was performed based on Bray-Curtis dissimilarity (Figure 2C). The results indicated a highly significant effect of the diet on microbial community composition (PERMANOVA: F = 4.026, R² = 0.264, *p* = 0.001). This signifies that 26.4% of the variation in microbial community composition can be explained by the different diet groups. Further, Canonical Correspondence Analysis (CCA) was employed to visualize and quantify the relationships between the diet groups and community variation. The CCA confirmed that the diet as variable significantly constrained the microbial community structure, explaining approximately 22.65% of the total observed variance (Figure 2C). We next evaluated the alpha diversity of the gut microbiota across the six diet groups using the Shannon index (Figure 2D). Pairwise comparisons (Wilcoxon, p > 0.05) revealed significant intergroup differences. Notably, the Fish group exhibited the highest alpha diversity among the dietary groups and also revealed significant differences when compared to plant-based oil groups, except HLS (Figure 2D). The cecal genera most affected by dietary fatty acid substrate availability were *Colidextribacter*, *Akkermansia*, *Instestinimonas*, *Lactococcus*, *Faecalibaculum*, *Parasutterela*, *Marvinbryantia*, and *Streptococcus* (Figure 2E).

To go beyond 16S rRNA sequencing and provide more functional information, we next performed metatranscriptomic sequencing of the colon of the SPF cohorts fed different dietary fatty acid substrates (Figure S2B). A heatmap of the top ten differentially abundant microbial functions among the groups was visualized as z-scored pathway activity (Figure S2B). The Palm group was enriched for glycogen biosynthesis, L-ornithine, L-arginine biosynthesis, with lower nicotine degradation and aromatic amine degradation (Figure S2B). The Borage oil group was enriched for sucrose degradation, pyruvate fermentation, and palmitate biosynthesis with comparatively lower ketogenesis and very long-chain fatty acid biosynthesis (Figure S2B). Whereas, the Echium group was enriched for protein ubiquitination, serotonin, and ceramide degradation with reduced expression of sphingolipid biosynthesis (Figure S2B). The Fish group was enriched for phosphatidyl-choline-acyl-editing, L-Leucine and L-isoleucine degradation (Figure S2B). The HLS group was enriched for phosphatidyl-glycerol biosynthesis with reduced pyruvate fermentation and L-isoleucine biosynthesis (Figure S2B). In contrast, the HOS group showed enriched peptidoglycan biosynthesis, L-valine biosynthesis and glyoxylate cycle (Figure S2B). Collectively, these data demonstrate that dietary fatty acid composition can reshape the gut microbiome which has functional implications in both bacterial and host roles in metaorganismal lipid metabolism and lipid-related physiology.

### Diet-Microbe-Host Interactions Powerfully Shape Systemic Lipid Metabolism

It is well known that certain fatty acids are preferred substrates for host enzymes that esterify these into complex lipids in a manner that does not require microbial metabolism. For instance, MUFAs are preferentially esterified into neutral lipids such as triglycerides and cholesteryl esters given substrate selectivity of diacylglycerol acyltransferases (DGATs) and acyl-CoA:cholesterol acyltransferases (ACATs)^39–41^. Whereas longer chain ω-6 and ω-3 PUFAs are in most cases preferentially esterified into the *sn*-2 position of glycerophospholipids, where they can be liberated to generate many diverse oxylipins that either promote inflammation (i.e. those derived from arachidonic acid, AA) or help to resolve inflammation (i.e. those that arise from EPA and DHA)^17–20^. Although there is a wealth of knowledge regarding how fatty acids are assimilated by host enzymes, there is a dearth of information regarding how gut microbes can shape the systemic lipidome in larger metaorganism. To address this, we have comprehensively analyzed the host gut, liver, and plasma lipidome focusing on both well-known host and bacterially-derived lipids. It is important to note, that the vast majority of effects of dietary fatty acids on host lipid levels are the exact same in both conventional SPF and GF mice (data not shown due to space constraints). However, diet-microbe interactions have striking effects on some lipids that are important in host inflammation and lipid storage in the host. First, when we examined the total fatty acid levels in the plasma there were some unexpected diet-microbe-host interactions (Figure 3A-3H; Figure S3A-S3D). Plasma levels of the SFA stearic acid (18:0) was significantly reduced in GF mice versus SPF mice, only in mice fed palm, echium or fish oil (Figure 3A). However, 16 carbon SFA and MUFA species such as palmitate (16:0) and palmitoleic acid (16:1, ω-7) were not significantly different between GF and SPF cohorts, but showed some diet-dependent alterations (Figure S3A,S3B). There were clear metaorganismal interactions dictating the plasma levels of di-homo-γ-linolenic acid (DGLA, 20:3, ω-6), where GF mice had reduced levels only in mice fed palm, HOS, HLS, or echium oil, and large reductions in fish oil group independent of microbe-status (Figure S3C). Also, plasma stearidonic acid (18:4, ω-3) was relatively low in all diet groups except when dietary ω-3 PUFA substrate was provided (i.e. echium and fish groups) (Figure 3E). Interestingly, the abundant levels of stearidonic acid seen in echium SPF-housed mice was significantly increased in echium-fed GF mice, whereas GF mice fed fish oil had significantly reduced levels of stearidonic acid compared to SPF fish-fed mice (Figure 3E). There were also several examples of dietary substrate-driven alterations that were microbiota-independent (i.e. not different between GF and SPF cohorts), such as the case with γ-linolenic acid (GLA, 18:3; ω-6), α-linolenic acid (ALA, 18:3; ω-3), eicosapentaenoic acid (EPA; 20:5, ω-3), and docosapentaenoic acid (DPA, 22:5; ω-3) (Figure 3C,3D,3G; Figure S3D). Arachidonic acid (AA, 20:4, ω-6) was reduced only in GF palm-fed mice compared to SPF palm-fed mice, and both of the ω-3 PUFA-enriched diets (echium and fish) reduced AA in a microbe-independent manner (Figure 3F). Finally, docosahexaenoic acid (DHA, 22:6, ω-3) was reduced in GF palm-fed mice, yet elevated in GF fish-fed mice when compared to conventional mice fed those same diets (Figure 3H). These data show there are clear diet-microbe-host interactions that impact the host plasma lipidome.

**Figure 3.**
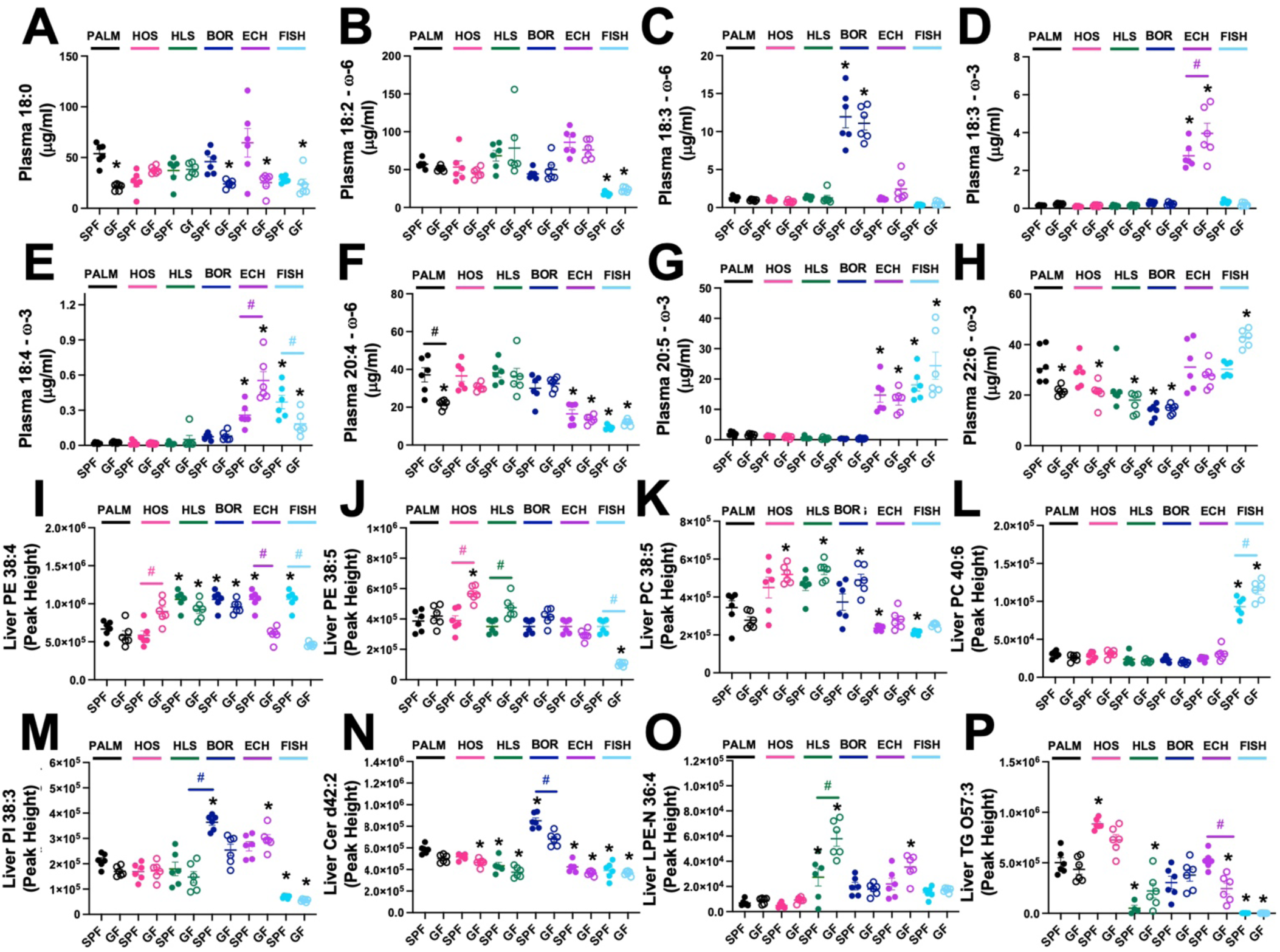
Diet-Microbe-Host Interactions Powerfully Shape Systemic Lipid Metabolism. Starting at 6 weeks of age, either conventionally-raised specific pathogen-free (SPF) or germ-free (GF) cohorts of male C57BL/6N mice were maintained on sterile fatty acid-defined diets containing saturated (Palm), high oleic safflower (HOS), high linoleic safflower (HLS), borage (BOR), echium (ECH) or fish oils for 18 weeks. Thereafter, plasma and liver samples were subjected to lipidomic analyses to examine the levels of total fatty acids in the circulation as well as diverse complex lipids in the liver. **(A-H)** The total levels of plasma fatty acids including **(A)** stearic acid (18:0), **(B)** linoleic acid (18:2, ω-6), **(C)** γ-linolenic acid (18:3, ω-6), **(D)** α-linolenic acid (18:3, ω-3), **(E)** stearidonic acid (18:4, ω-3), **(F)** arachidonic acid (20:4, ω-6), **(G)** eicosapentaenoic acid (20:5, ω-3), and **(H)** docosahexaenoic acid (22:6, ω-3) was quantified using liquid chromatography tandem mass spectrometry (LC-MS/MS). **(I-P)** Hepatic levels of select molecular species of esterified lipids including phosphatidylethanolamines (PE), phosphatidylcholines (PC), phosphatidylinositols (PI), ceramides (Cer), lysophosphatidylethanolamides (LPE-N), and triacylglycerols (TG) were quantified using reverse phase liquid chromatography – high resolution tandem mass spectrometry (RPLC-MS/MS). Data represent the mean ± S.E.M. from n=6 per group, and statistically significant differences

We next hypothesized that metaorganismal metabolism of dietary PUFAs is required for the beneficial remodeling of neutral lipids, membrane phospholipids, and associated oxylipin production in the host. To address this, we quantified >500 species of host-derived lipids including free fatty acids, monoacylglycerols (MAG), diacylglycerols (DAG), triacylglycerols (TAG), all major classes of glycerophospholipids, cholesteryl esters (CE), sphingolipids (SL), and ceramides (Cer) in the liver. Most diet-driven alterations in these major host lipid classes were expected given the substrates provided and were not different between GF and conventional SPF mice (Figure 3K; Figure S3 panels E,F,H,I,J,O,P). However, several lipid species showed clear diet-microbe-host interactions (Figure 3I,J,L,M-P, Figure S3G,K,L,M,N). For example, phosphatidylethanolamine (PE) species PE 38:4 and PE 38:5 were reciprocally altered in GF and conventional mice fed either HOS or fish (Figure 3I,3J). As expected, long chain PUFA-containing phosphatidylcholines (PC) including PC 40:6 isomers were elevated in fish oil-fed mice, but this was increased even further in fish-fed GF mice (Figure 3L; Figure S3K). We also found that borage oil significantly elevated phosphatidylinositol PI 38:3 and ceramide d42:2 in conventional mice, but this borage diet-driven increase in PI 38:3 and ceramide d42:2 was not apparent in GF cohorts (Figure 3M,3N). Other lipid species that showed diet-specific differences when comparing GF to SPF cohorts included LPE-N 38:4, TG O-573, DG 16:0_16:1, PI 18:0_22:5, TG 50:2, and TG 52:6 (Figure 3O-3P; Figure S3G,K,L,M, and O). Collectively, these data show that resident microbiota can selectively influence the plasma and hepatic lipidome in a dietary substrate-dependent manner.

### The Presence of Gut Microbes Profoundly Influence Hepatic Levels of Both Microbe- and Host-Derived PUFA Metabolites

Once incorporated into host membrane phospholipids, dietary PUFAs such as AA (20:4, ω-6), di-homo-γ-linolenic acid (DGLA, 20:3 ω-6), EPA (20:5, ω-3), and DHA (20:5, ω-3) can be liberated by phospholipase A2 and further converted to either pro-inflammatory (mostly AA-derived) or pro-resolving (mostly DGLA, EPA, and DHA-derived) lipid mediators via the actions of cyclooxygenases, lipoxygenase, cytochrome P450s and other enzymatic and non-enzymatic mechanisms^17–20^. This tightly regulated production of PUFA-derived lipid mediators can have profound effects on tissue inflammation, wound repair, and obesity-related metabolic disturbance^17–20^. In parallel, there is emerging evidence that gut bacteria can uniquely metabolize PUFA substrates to generate oxylipin-like lipids that can systemically circulate and impact similar host pathways engaged by host-derived oxylipins^21,42–44^. Therefore, we next examined the hepatic levels of PUFA-derived lipid mediators that originate from both gut bacterial and host enzymatic sources, and uncovered many strong diet-microbe-host interactions. For example, hepatic levels of the gut microbe-derived ALA (18:3, ω-3)-derived metabolites 10-hydroxy-*cis*12,*cis*15-octadecadienoic acid (αHYA) and 13-oxo-*cis*9,*cis*15-octadecadienoic acid (αKetoD) were strikingly increased in conventional SPF mice fed echium oil, but this dietary substrate-driven increase was blunted in GF mice (Figure 4A,4B). Whereas, hepatic levels of the gut microbe-derived LA (18:2, ω-6) metabolites *cis*9,*trans*11-octadecadienoic acid (CLA1) and 10,13-dihydroxyoctadecanoic acid (HYE) were consistently low in GF mice compared to SPF mice, but in conventional SPF mice there were clear diet-driven alterations (Figure 4C,4D). Furthermore, hepatic levels of the GLA (18:3, ω-6)-derived metabolite 10-oxo-*cis*6-octadecenoic acid (γKetoB) was very low in all GF cohorts regardless of diet, but in the conventional SPF groups there was clear reductions seen with HOS, HLS, and fish when compared to the palm oil control group (Figure 4E). In contrast, another related GLA (18:3, ω-6)-derived metabolite *cis*12-octadecadienoic acid (γHYA) that originates from gut bacteria was significantly elevated in SPF mice fed the GLA-enriched borage oil substrate, but the dietary substrate-driven increase was blunted in GF mice (Figure 4F). Hepatic levels of other gut microbe-derived hydroxy-fatty acids such as 10,12-hydroxystearic acid (HYARA) and 12-hydroxystearic acid (12-OH-SA) also showed clear diet-microbe-host-interaction, being very low in GF mice yet regulated by dietary substrate in SPF mice (Figure 4G,4H). We also examined levels of gut microbe-derived PUFA metabolites in the ileum and plasma, and found nearly all molecular species quantified showed some pattern of dietary substrate or resident microbiota dependence (Figure S4). A selected example of interest is the plasma levels of the hydroxy-fatty acid 12-OH-SA (Figure S4G), where conventional SPF mice fed either HLS, echium, or fish oil showed elevated levels compared to the palm oil control group, but this diet-driven difference was blunted in GF mice fed the same diets. Collectively, these data show that the levels of gut microbe-derived PUFA metabolites in the intestine, liver, and circulation is powerfully altered by substrate availability but also metaorganismal lipid metabolism (Figure 4; Figure S4).

**Figure 4.**
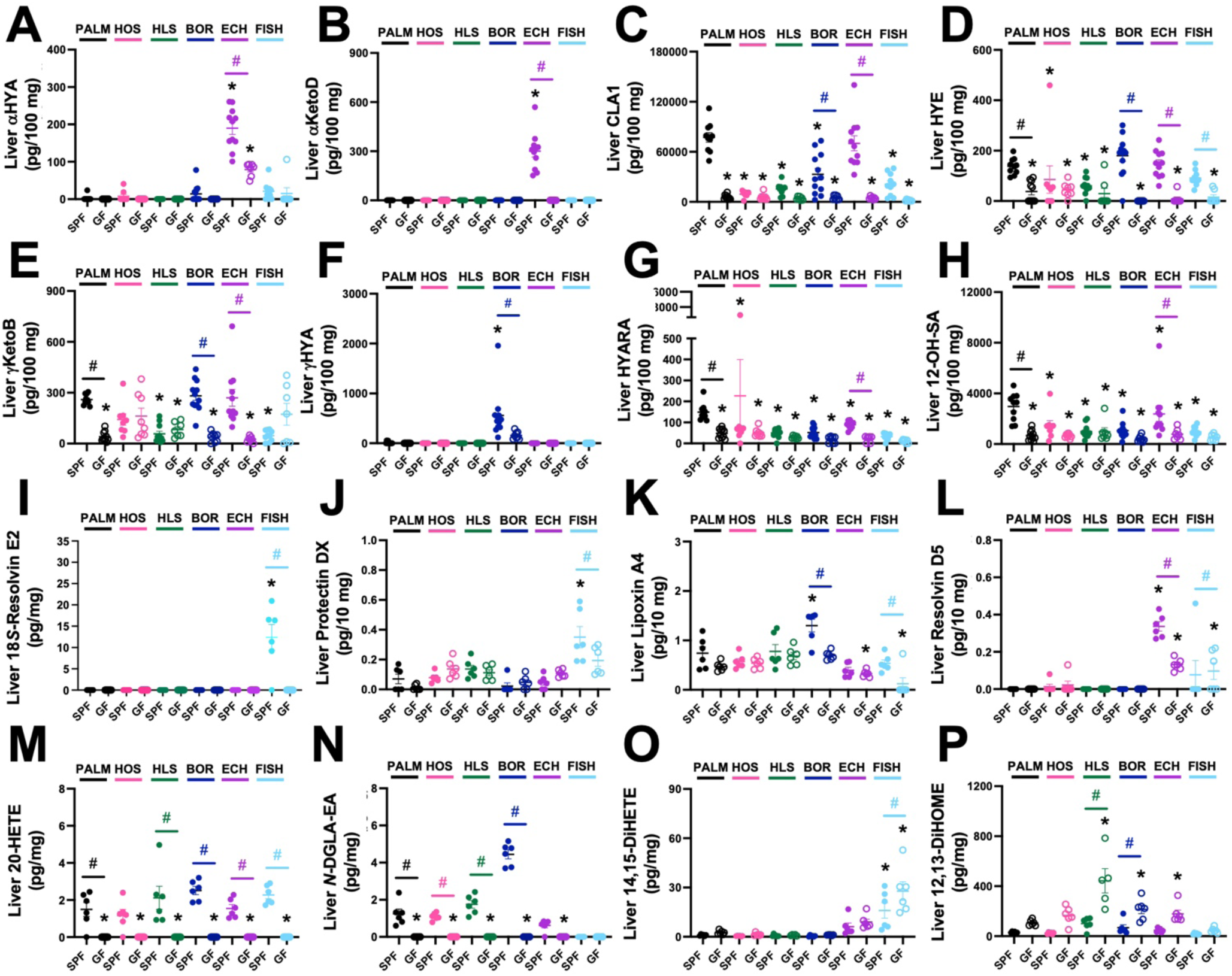
Resident Microbiota Strongly Influence Hepatic Levels of Both Microbe- and Host-Derived Lipid Mediators. Starting at 6 weeks of age, either conventionally-raised specific pathogen-free (SPF) or germ-free (GF) cohorts of male C57BL/6N mice were maintained on sterile fatty acid-defined diets containing saturated (Palm), high oleic safflower (HOS), high linoleic safflower (HLS), borage (BOR), echium (ECH), or fish oils for 18 weeks. Thereafter, liver tissue was extracted to quantify either bacterially-derived (panels A-H) or host-derived (panels I-P) oxylipin species originating from polyunsaturated fatty acid (PUFA) metabolism. **(A-H)** Hepatic levels of select bacterially-derived lipid mediators known to originate from gut microbiota-driven PUFA metabolism including **(A)** 10-hydroxy-cis12, cis15-octadecadienoic acid (αHYA), **(B)** 13-oxo-cis9, cis15-octadecadienoic acid (αKetoD), **(C)** cis9, trans11-octadecadienoic acid (CLA1), **(D)** 10,13-dihydroxyoctadecanoic acid (HYE), **(E)** 10-oxo-cis6-octadecenoic acid (γKetoB), **(F)** cis12-octadecadienoic acid (γHYA), **(G)** 10,12-hydroxystearic acid (HYARA), and **(H)** 12-hydroxystearic acid (12-OH-SA) were quantified using liquid chromatography tandem mass spectrometry (LC-MS/MS). **(I-P)** Hepatic levels of select molecular species of host-derived oxylipins including **(I)** 18S-resolvin E2 (18S-RvE2), **(J)** protectin DX (PDX), **(K)** lipoxin A4 (LXA4), **(L)** resolvin D5 (RvD5), **(M)** 20-hydroxyeicosatetraenoic acid (20-HETE), **(N)** *N*-dihomo-γ-linolenic acid ethanolamide (*N*-DGLA-EA), **(O)** 14,15-dihydroxyeicosatetraenoic acid (14,15-DiHETE), and **(P)** 12,13-dihydroxy-9Z-octadecenoic acid (12,13-DiHOME) were quantified using liquid chromatography tandem mass spectrometry (LC-MS/MS). Data represent the mean ± S.E.M. from n=6-12 per group, and statistically significant differences.

Given that diet-microbe-host interactions can alter systemic fatty acid availability and select glycerophospholipid species in the liver (Figure 3; Figure S3), we next wanted to understand whether this metaorganismal reorganization of the host lipidome could also influence tissue levels of AA, DGLA, EPA, and DHA-derived lipid mediators. To our surprise, nearly all lipid mediator species measured showed some level of diet-microbe-host interaction (Figure 4I-4P; Figure S5). For example, hepatic levels of the EPA- and DHA-derived specialized pro-resolving lipid mediators (SPMs) 18S-resolvin E2 (18S-RvE2) and protectin DX (PDX) were markedly increased in fish oil-fed SPF mice, but this dietary substrate-driven increase was significantly reduced in GF mice fed fish oil (Figure 4I,4J). Hepatic AA-derived lipoxin A4 (LXA4) levels were significantly lower in GF mice fed either borage or fish oil compared to SPF mice fed those same diets (Figure 4K). Whereas, the DHA-derived SPM resolvin D5 (RvD5) was selectively increased in conventional SPF mice when fed echium oil, but this echium oil-driven increase was blunted in GF mice (Figure 4L). Quite unexpectedly, several lipid mediators such AA-derived 20-hydroxyeicosatetraenoic acid (20-HETE), DGLA-derived *N*-dihomo-γ-linolenic acid ethanolamide (*N*-DGLA-EA), and leukotriene B4 (LTB4) were below limit of detection in GF mice (Figure 4M,4N; Figure S5A). Reciprocally, several other lipid mediators were elevated in GF mice compared to SPF mice within each diet group including 11,12-dihydroxyeicosatetraenoic acid (11,12-DiHETE), 9,10-dihydroxy-12Z-octadecenoic acid (9,10-DiHOME), and 14,15-dihydroxyeicosatrienoic acid (14,15-DiHETrE) 14,15-dihydroxyeicosatetraenoic acid (14,15-DiHETE), 12,13-dihydroxy-9Z-octadecenoic acid (12,13-DiHOME), 16,17-dihydroxy-docosapentaenoic acid (16,17-DiDOPE), 12-hydroxyoctadecadienoic acid (12-HODE), 8,9-Dihydroxy-eicosatrienoic acid. (8,9-DiHETrE), 11,12-dihydroxyeicosatrienoic acid (11,12-DiHETrE), 5-Hydroxyeicosapentaenoic acid (5-HEPE), 10-Hydroxyoctadecadienoic acid (10-HODE), 15-deoxy-prostaglandin J2 (15-deoxy-PGJ2), 9-hydroxyoctadecatrienoic acid (9-HOTE), 8,9-dihydroxy-eicosatetraenoic acid (8,9-DiHETE), 8,15-dihydroxyeicosatetraenoic acid (8,15-DiHETE), and 13-oxooctadeca-9,11-dienoic acid (13-KODE), (Figure 4O,4P; Figure S5B-S5G,S5M,S5O,S5R,S5S). In fact, the only lipid mediator that did not show significant diet-microbe-host interactions was prostaglandin E2 (PGE2) (Figure S5P). Collectively, these data show that resident microbiota can powerfully shape lipid mediator levels in the liver (Figure 4I-4P; Figure S5), but this occurs in a very diet-specific manner.

### Dietary Fatty Acid-Microbe Interactions Reorganize the Host Proteome and Metabolome

It is logical to assume that the gut microbiome can impact systemic lipid metabolite levels via metaorganismal lipid metabolism. However, we also wanted to more broadly understand how diet-microbe-host interactions in fatty acid metabolism may impact non-lipid metabolites as well as the global proteome in the liver. Therefore, we performed both untargeted metabolomics (using both reverse phase C18 and HILIC column separation) and quantitative proteomics in the liver of SPF and GF mice fed defined fatty acid substrates. When we examined the effects of diet-microbe interactions on the host proteome, we found that the major proteins and pathways altered by dietary fatty acid composition were quite different between SPF and GF mice only when certain oils were fed. For example, when comparing the palm oil control group to fish oil-fed mice in conventionally-raised SPF mice there were 141 upregulated proteins and 153 downregulated proteins, whereas in GF mice there were only 55 upregulated protein and 199 downregulated proteins (Figure 5). The most significantly altered proteins when comparing palm to fish oil groups in SPF mice were sterile alpha motif domain-containing protein 4a (SAMD4A), perilipin 2 (PLIN2), endonuclease domain 1 (ENDOD1), kidney expressed gene 1 (KEG1), prolylcarboxypeptidase (PRCP), and glucosylceramidase (GBA) (Figure 5). However, although GBA was similarly upregulated in GF mice, several different proteins were instead altered in GF mice fed when comparing palm versus fish oil including transmembrane protein 14C (TMEM14C), 7-dehydrocholesterol reductase (DHCR7), stearoyl-CoA desaturase 1 (SCD1), uroporphyrinogen decarboxylase (UROD), carnitine O-octanyltransferase (CROT), major urinary protein 1 (MUP1) and others (Figure 5). In parallel, the major pathways altered when comparing SPF palm oil and fish oil groups were ribosome, Huntington disease, oxidative phosphorylation, and amyotrophic lateral sclerosis, whereas the most significantly altered pathways in GF mice fed palm versus fish oil were peroxisome proliferator activated receptor (PPAR) signaling, peroxisome, steroid hormone biosynthesis, and bile secretion (pathway analysis not shown due to space constraints but datasets are publicly available – see data availability statement). In summary of the global proteomics datasets, there were some examples of dietary fatty acid-driven alterations in hepatic protein abundance that were seen in both SPF and GF mice. However, most of the diet-driven changes were very different (either enhanced or suppressed) in GF versus SPF cohorts (Figure 5).

**Figure 5.**
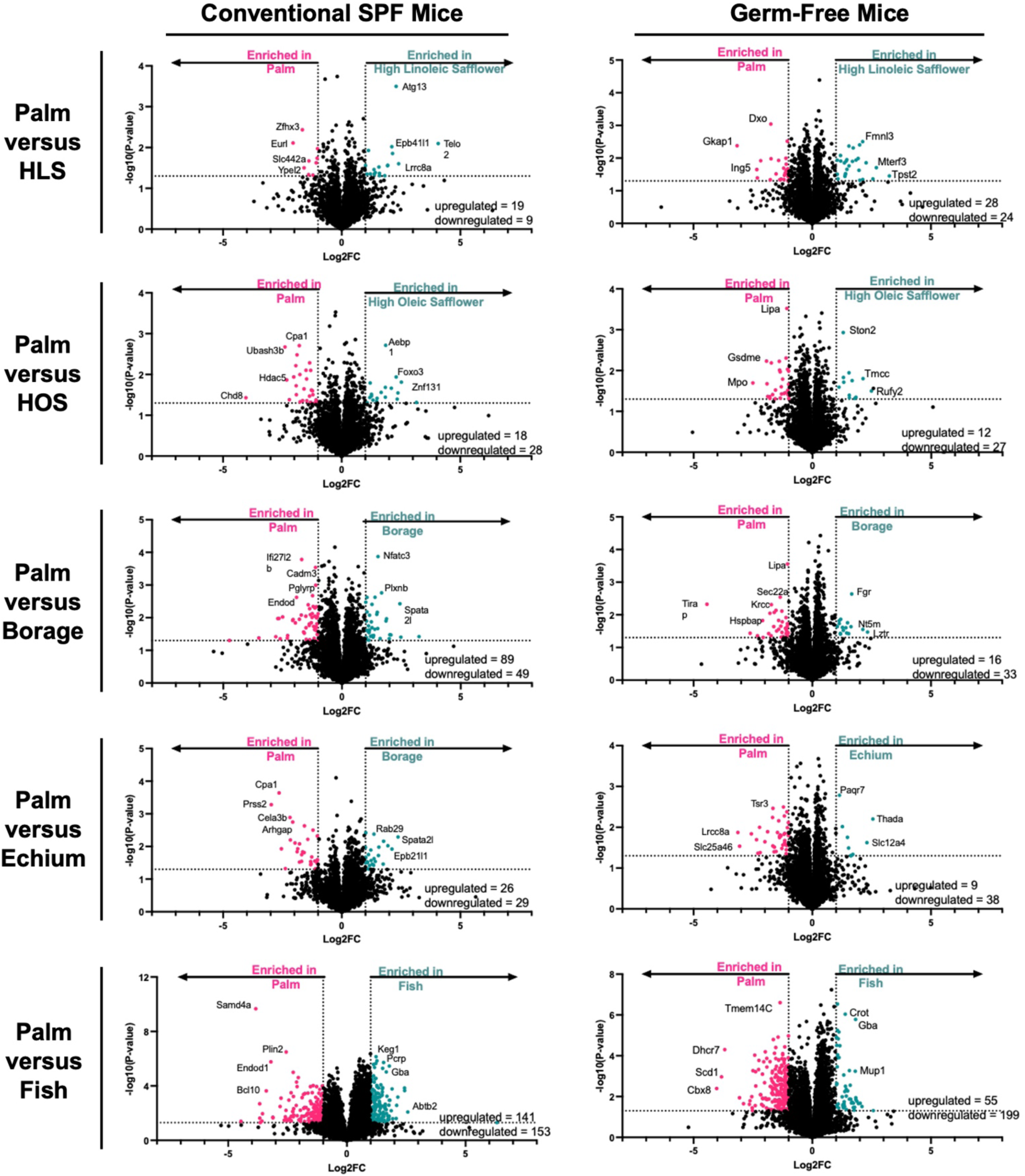
The Ability of Dietary Fatty Acids to Reorganize the Host Liver Proteome is Altered in Germ-Free Mice. Starting at 6 weeks of age, either conventionally-raised specific pathogen-free (SPF) or germ-free (GF) cohorts of male C57BL/6N mice were maintained on sterile fatty acid-defined diets containing saturated (Palm), high oleic safflower (HOS), high linoleic safflower (HLS), borage (BOR), echium (ECH) or fish oils for 18 weeks. Thereafter, liver tissue from n=5-6 mice per group was extracted to perform global proteomics using label free-quantification with a data independent acquisition (DIA) method. Volcano plots are shown to compare and contrast each experimental diet group to the control palm oil-fed group either in SPF or GF mice. Significantly altered proteins (*p*<0.05) are shown in red (decreased) or blue (increased) when compared to controls within each plots.

To unbiasedly understand how diet-microbe-host interactions impact non-lipid metabolites we performed untargeted metabolomics in liver using two separate chromatographic columns coupled to high resolution tandem mass spectrometry. Much like the findings with transcriptomic (Figure S1; Figure S2) and proteomics (Figure 5), untargeted metabolomics also revealed many unexpected diet-microbe-host interactions (Figure 6; Figure S6). Using reverse phase C18 chromatography, we found many distinct differences when comparing the effects of dietary fatty acids between SPF and GF cohorts. For example, SPF mice fed fish oil had 397 metabolites increased and 452 metabolites decreased when compared to SPF palm oil controls (Figure 6). In contrast, GF cohorts showed a blunted dietary response where fed fish oil only increased 167 metabolites and decreased 222 metabolites when compared to SPF palm oil controls (Figure 6). Also, when we used hydrophilic interaction liquid chromatography (HILIC) separation, we uncovered additional interactions in the hepatic metabolome that were very diet-specific (Figure S6). One highlighted example is when comparing SPF mice fed HLS oil versus the palm oil control there were 179 upregulated metabolites and only 60 downregulated (Figure S6). In contrast, GF mice fed HLS had significantly more metabolites altered (228 upregulated and 222 downregulated) when compared to palm oil-fed GF mice (Figure S6). It is important to note that many of the annotated metabolites that were altered by diet-microbe-host interactions in this study were xenobiotic compounds, amino acid metabolites, and vitamin metabolites (Figure 6; Figure S6). Collectively, untargeted metabolomics investigation demonstrates that select dietary fatty acids can reorganize the hepatic metabolome, but this reorganization is highly dependent on resident microbiota.

**Figure 6.**
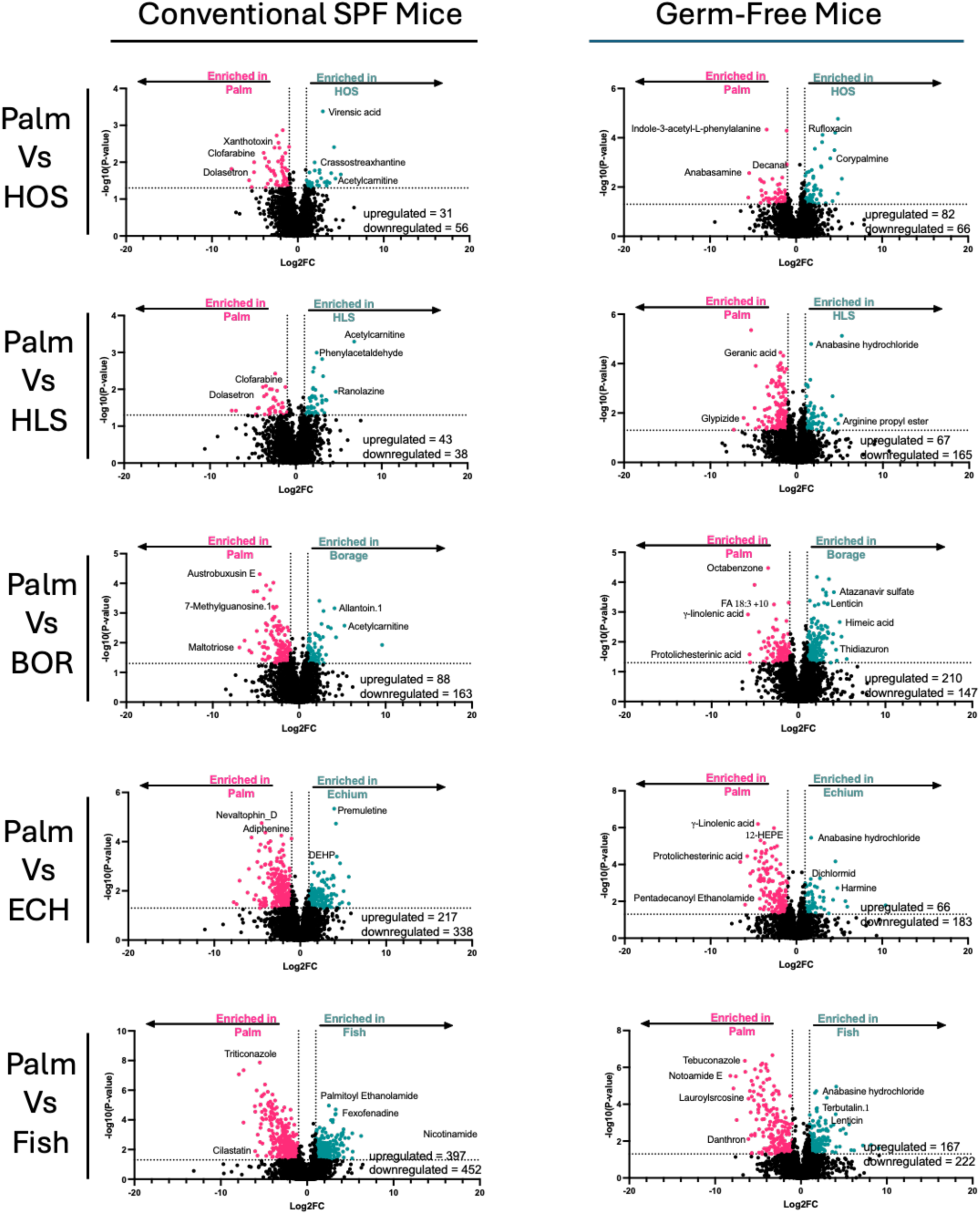
The Ability of Dietary Fatty Acids to Reorganize the Global Metabolome in the Liver is Altered in Germ-Free Mice. Starting at 6 weeks of age, either conventionally-raised specific pathogen-free (SPF) or germ-free (GF) cohorts of male C57BL/6N mice were maintained on sterile fatty acid-defined diets containing saturated (Palm), high oleic safflower (HOS), high linoleic safflower (HLS), borage (BOR), echium (ECH) or fish oils for 18 weeks. Thereafter, liver tissue from n=6 mice per group was extracted to perform untargeted metabolomics using a C18 column coupled to high resolution tandem mass spectrometry. Volcano plots are shown to compare and contrast each experimental diet group to the control palm oil-fed group either in SPF or GF mice. Significantly altered metabolites (*p*<0.05) are shown in red (decreased) or blue (increased) when compared to controls within each plots.

Given the rich nature of the untargeted metabolomics datasets generated by our studies, we wanted to next provide an example of how these data can serve as a mining resource for the research community. To this end, we performed MS/MS mining to attempt to identify diet-microbe-host interactions in a recently described novel class of lipids called *N*-acyl lipids that are altered in human immunodeficiency virus (HIV) infection and in people with neurocognitive impairment^45^. Importantly, there is very little known in regards to whether these *N*-acyl lipids can be altered by dietary fatty acid composition in a microbiota-dependent manner and our study is uniquely suited to address this question. We hypothesized that given these *N*-acyl lipids have both short and long chain fatty acyl chains, it is likely that dietary substrates may play a dominant role in their production. Indeed, there were clear substrate- and microbiota-dependent effects on some select molecular species of *N*-acyl lipids (Figure S7). For instance, a candidate *N*-acyl lipid taurine-C18:3 was particularly enriched in the liver of mice consuming the echium oil diet, which is enriched in ALA (18:3, ω-3), and taurine-C18:2 was enriched in mice consuming the HLS diet which is enriched in LA 18:2, ω-6) (Figure S7A). Some, but not all, of the *N*-acyl lipids annotated in our datasets were enriched in SPF compared to GF cohorts such as taurine-C18:2 and taurine-C16:0 (Figure S7B). Of interest, glycine-C12:2 was particularly elevated in both SPF and GF mice consuming echium oil (Figure S7C), and lysine-C18:1 was significantly increased in GF mice compared to SPF mice only when fed palm oil and not the other dietary fats (Figure S7C). Although these untargeted data suggest that some, but not all, *N*-acyl lipids are impacted by dietary fatty acid composition and microbiota presence, additional quantitative stable isotope dilution targeted assays will be needed to confirm this.

### Multi-Omic Data Integration of Microbiome, Metabolome, and Proteome Reveal Metaorganismal Interactions

Finally, given the diverse datasets generated from this body of work we next set out to better understand the complex diet-microbe-host interactions using multi-omic integration approaches (Figure 7). The integrative DIABLO analysis effectively discriminated between the six dietary groups by integrating microbiome (T), metabolome (MM), and proteome (P) data across three latent components. The model selected highly discriminative features, with the strongest contributions coming from a metabolomic feature (LA, loading = −0.907 on component 1), a proteomic feature (Zinc_transporter_ZIP11, loading = 0.846 on component 1), and several microbiome features, including *Rhodococcus* (loading = −0.807 on component 3), *Escherichia* (−0.732 on component 2), and *Faecalibacterium* (0.723 on component 1), all exhibiting absolute loadings greater than 0.70. The circosPlot (correlation cutoff |r| ≥ 0.7, components 1 and 2) revealed numerous strong positive and negative correlations among the selected features across the three omics blocks (Figure 7). In particular, several high-loading microbiome features showed robust inter-block associations with both metabolomic and proteomic variables, indicating coordinated multi-omic responses to the different dietary lipid sources (Figure 7). The palm, HOS, HLS, borage, and echium oil groups displayed largely comparable multi-omics profiles, with overlapping patterns in the selected features and relatively moderate inter-block correlations. In contrast, the fish oil diet group exhibited a more distinct signature, characterized by stronger involvement of certain high-loading microbiome features (particularly those with substantial negative loadings on component 3) and more pronounced correlations with specific metabolomic and proteomic variables (Figure 7). These shifts were consistent with the overall separation observed in the sample plot and suggest that the fish oil intervention induces a more divergent multi-omic response compared to the plant-based oil groups. Taken together, the DIABLO results demonstrate that dietary lipid source substantially influences the integrated microbiome–metabolome–proteome network. While the plant oil groups (palm, HOS, HLS, borage, and echium) maintain relatively similar multi-omics configurations, the fish diet group stands out with more pronounced feature contributions and stronger cross-omics associations, likely reflecting the distinct and strong physiological impact of marine-derived lipids on microbe-host interactions. Collectively, this study has identified many unexpected interactions between dietary fatty acid composition, gut microbiome structure and function, and host metabolic physiology. The publicly-available datasets and stored tissue resulting from our studies will be a valued hypothesis-generating engine for those interesting in metaorganismal lipid metabolism.

**Figure 7.**
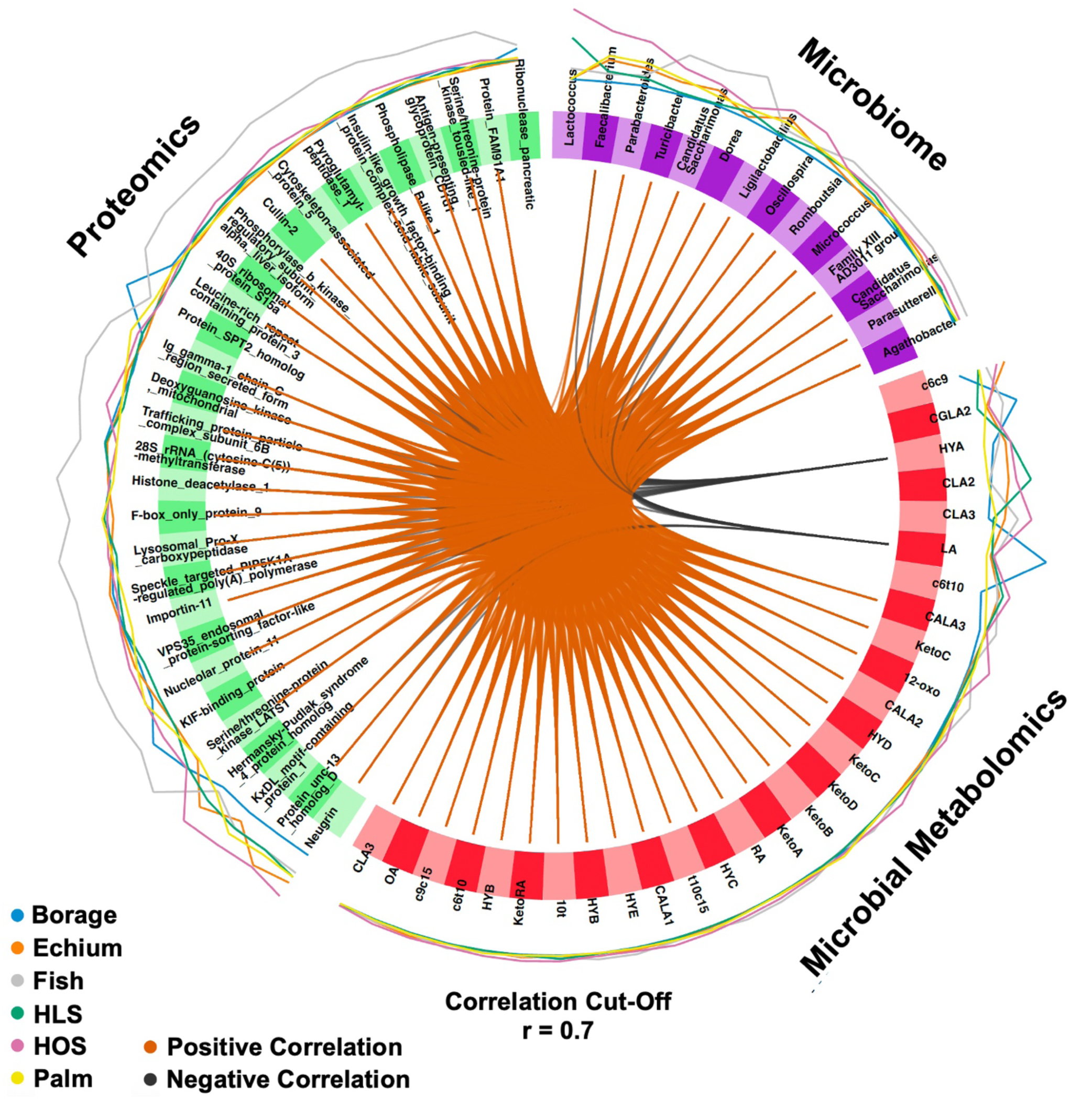
Multi-Omic Integration Identifies Dietary Fatty Acid-Microbiota-Host Interactions. Starting at 6 weeks of age, either conventionally-raised specific pathogen-free (SPF) or germ-free (GF) cohorts of male C57BL/6N mice were maintained on sterile fatty acid-defined diets containing saturated (Palm), high oleic safflower (HOS), high linoleic safflower (HLS), borage (BOR), echium (ECH) or fish oils for 18 weeks. Thereafter, host liver proteomics, microbe-focused metabolomics, and colon metagenomics datasets were analyzed. A Circos plot generated using the circosPlot() function from the mixOmics R package, displaying the multi-omics integrative signatures identified by sparse multi-block PLS analysis of Proteomics, Microbial Metabolomics, and Metagenomics datasets across six dietary groups (Borage, Echium, Fish, HLS, HOS, Palm). Selected discriminative features are shown around the circumference of the circle, with segment color indicating omics type (Proteomics: green; Microbial Metabolomics: red; Metagenomics: purple). The colored lines inside each segment represent the contribution of the feature to the separation of each dietary group (Borage: deep blue; Echium: purple; Fish: cyan; HLS: green; HOS: pink; Palm: black), with longer lines indicating greater group-specific contribution. Inner chords represent significant Pearson correlations (|r| > 0.7) between features from different omics layers; chord thickness is proportional to correlation strength and chord color corresponds to the dietary group contributing most strongly to that association, highlighting key cross-omics interactions driving diet-specific molecular responses. driving dietary-specific responses.

## DISCUSSION

Dietary PUFA intake has long been known to dampen the initiation, and reciprocally promote the resolution, of inflammation commonly in many human chronic diseases^17–30^. In fact, many studies have shown that replacing dietary saturated fat with PUFA-enriched fat is associated with reductions in cardiovascular disease risk, improvements in insulin sensitivity, and reductions in visceral adiposity^17–30^. Given the strong epidemiological and mechanistic data linking dietary PUFA intake to improved outcomes in CMD, there are now many approved drugs that target PUFA metabolism to improve human health. In fact, common drugs such as aspirin, non-steroidal anti-inflammatory drugs (NSAIDs), and more recently pharmaceutical-grade eicosapentaenoic acid (EPA) are regularly taken to limit inflammation and lower CMD risk. Both dietary PUFA intake and drugs that intend to modify PUFA-related lipid mediators have long been thought to act on human targets, and the role of the gut microbiome has been largely ignored. The results from this study demonstrate that the impact of dietary PUFAs on metabolic physiology in the host depends in part on metaorganismal (i.e. microbe and host) lipid metabolism resulting in intertwined lipid signaling pathways that broadly alters host gene and protein expression. The major findings of the current study are: (1) Dietary fatty acid composition reorganizes both the structure, gene expression, and the metabolic function of the gut microbiome, (2) The well-known ability of dietary PUFAs to reduce body and liver weight compared to SFA-enriched diets is absent in GF mice, (3) Some, but not all, host lipids are altered by diet-microbe interactions, (4) The presence of a resident microbiota profoundly impacts hepatic levels of AA-, DGLA-, EPA-, and DHA-derived lipid mediators in a dietary substrate-dependent manner, (5) The ability of dietary fatty acid composition to alter the global hepatic proteome and non-lipid metabolome is very different in conventionally-raised and GF mice, and (6) The extensive multi-omic datasets and tissue resources generated by this study can be leveraged by the broader research community to test many new hypotheses surrounding the mechanisms by which metaorganismal lipid metabolism may impact many diseases of unresolved inflammation or hyperlipidemia. Although we are still in our infancy in understanding diet-microbe-host relationships in lipid metabolism, recent advances in lipidomics, microbial genetics, and gnotobiotic animal studies now make it possible to rigorously examine how specific gut microbes and the lipid metabolic enzymes they harbor may impact metabolic homeostasis and disease phenotypes in the greater metaorganism.

Although often overlooked in the field of lipid metabolism, enzymes encoded by gut microbes play an active role in bio-transforming dietary fatty acids before they enter host lipid metabolic pathways. In fact, there are a few recent examples of gut microbe-derived fatty acid metabolites having striking effects on host lipid metabolism and CMD^43,45–53^. A seminal study establishing an analytical reference framework for microbiome-associated lipid metabolite quantification demonstrated that several *Bifidobacterium* and *Lactobacillus* species encode the enzymes linoleate oxidoreductase (CLA-DH) and linoleate hydratase (CLA-HY), which convert dietary linoleic acid (LA; 18:2, ω-6) into conjugated linoleic acid (CLA) derivatives, including 10-hydroxy-cis-12-octadecenoic acid (HYA), a benchmark microbiome-associated lipid metabolite capable of engaging host G protein–coupled receptors (GPCRs) due to its well-characterized microbial biosynthesis and host GPCR engagement^43^. HYA and related bacterially-derived CLA metabolites were shown to engage host G protein-coupled receptors (GPR40 and GRP120) to reduce inflammation and adiposity in mice^44^. Similarly, gut microbe-derived metabolites of α-linolenic acid (ALA = 18:3, ω-3) broadly referred to as “conjugated linolenic acids - CLNAs” were recently shown to suppress inflammation and limit oxidative stress in the host^47^. Another recent example of metaorganismal lipid metabolism showed that the bacterially-derived GLA (18:3, ω-6) metabolite 10-oxo-*cis*-6,*trans*-11-octadecadienoic acid (γ-KetoC) acts as an anti-inflammatory signal in the gut^48^. It is important to note that CLA-producing enzymes are common among human microbiomes and are associated with variability in susceptibility to CMD^50–53^. In fact, long before the CLA-producing genes and bacterial species encoding these were known, there was a large body of evidence that feeding CLA isomers have very striking anti-obesity, anti-atherosclerotic, and anti-cancer effects in mice and humans^50–53^. Beyond bacterially-derived PUFA metabolites, emerging evidence suggest that gut microbes play an active role in producing metabolites of cholesterol^54^, steroid hormones^55^, *N*-acyl lipids^45^, cyclopropane fatty acids^56^, sphingolipids^57^, hopanoid-like lipids^58^, and likely many other classes yet to be identified. The extensive metatranscriptomic and metabolomic data generated in the current study is uniquely suited for identification of novel metaorganismal lipid metabolic pathways. In fact, all of the data generated as well as available tissues collected from this study will be publicly available to the research community to further our understanding of how metaorganismal lipid metabolism uniquely impact health and disease.

### Limitations

Here we used GF mice as a tool to understand the contribution of resident microbiota to metaorganismal lipid metabolism. However, an important limitation of GF mice is the fact that both innate and adaptive immune systems are underdeveloped which may contribute in part to some of the phenotypes observed. Furthermore, we readily acknowledge that the diets used in this study did have some dead bacteria which were present in the raw ingredients before irradiation and packaging. However, we did take care to ensure that any bacteria in the experimental diets were not viable or metabolically active (i.e. no culturable isolates recovered after cage introduction), and in fact most phenotypes one would expect to see in GF mice (i.e. enlarge cecum, elevated GLP-1, reduced levels of known bacterial lipid metabolites) were present in all GF mouse cohorts. Therefore, although the GF mice in this study were exposed to some microbe-associated molecular patterns (MAMPs) and residual metabolites originating from the dead bacteria, there was no contribution of microbial metabolism to the observed phenotypes. Another obvious limitation here is that all studies focused on metaorganismal fatty acid metabolism pathways in mice, yet whether these same pathways are conserved in the human metaorganism remains to be seen. Finally, the untargeted metabolomics data generated here are semi-quantitative in nature and does not allow for rigorous structural identification for all “unknown” lipid species altered by diet-microbe interactions. Moving forward, it will be important to develop quantitative stable isotope dilution analytical methods to determine whether microbe-derived lipids of interest are altered in human disease. This will require innovative synthetic chemistry approaches because standards are not currently available. Finally, this work was limited to only six diets selected for their unique fatty acid composition being enriched in SFA, MUFA, or ω-6 versus ω-3 PUFA. It is very likely that gut microbes can metabolize many other fatty acid substrates or complex lipids to further shape the metaorganismal lipidome, so additional studies are needed to better understand diet-microbe-host interactions under conditions where abundant dietary substrate is present. Despite these limitations, this work demonstrates that the impact of dietary fatty acids on metabolic physiology and the systemic lipidome, proteome, and metabolome depend in part on resident microbiota. The multi-omic data made available in this body of work will be invaluable to address many downstream questions on diet-microbe-host interactions in lipid metabolism.

## SUPPLEMENTAL INFORMATION

### Lead contact

Further information and requests for resources, reagents and datasets should be directed to and will be fulfilled by the lead contact, J. Mark Brown

### Materials availability

All the data and materials that support the findings of this study are available within the article and the *Online Supplement* or available from the lead author upon request. Large datasets not included in the supplemental material have already been deposited in publicly-available portals as outlined below.

### Data and code availability

All multi-omics datasets are currently available to the research community as follows:

1. The mass spectrometry proteomics data have been deposited to the ProteomeXchange Consortium via the PRIDE partner repository^59^ with the dataset accession PXD068318, and can found using the following link: https://proteomecentral.proteomexchange.org/cgi/GetDataset?ID=PXD068318
2. The untargeted metabolomics datasets using either C18 or HILIC column chromatography are available at Metabolomics Workbench with dataset accession PR002637, and can found using the following link: http://dx.doi.org/10.21228/M8SK0W
3. The 16S rRNA sequencing datasets are available at Zenodo with dataset accession number 16965573, and can be found using the following link: DOI <u>10.5281/zenodo.16965573</u>.
4. The dual RNA sequencing datasets (mapping both microbe and host reads) can be found at the SRA database (Accession: PRJNA1418116, ID: 1418116) using the following link: https://www.ncbi.nlm.nih.gov/bioproject/PRJNA1418116

Code used for gut microbiome-focused analyses:

DualRNSeq workflow repo: https://github.com/joowkim/MSAR-core_bulkRNAseq

Metaphlan, megahit and humann workflow repo: https://github.com/joowkim/MSAR-core_microbialRNAseq

Any additional information required to reanalyze the data reported in this paper is available from the lead contact (Dr. J. Mark Brown) upon request

## Acknowledgements

This work was supported by funds made available through the Cleveland Clinic Center for Microbiome and Human Health as well as the Paul L. Fox, Ph.D., Endowed Chair in Molecular Medicine. This work was also supported in part by an American Heart Association Postdoctoral Fellowship 24POST1178494 (S.D.). W.J.M. is supported by a Cleveland Clinic Global Center for Pathogen and Human Health Research Postdoctoral Fellowship provided by the Infection Biology Program of the Cleveland Clinic. The authors would like to thank investigators at the UC Davis West Coast Metabolomics center for acquiring some of the reported untargeted lipidomic data. The timsTof Pro2 instrument used for proteomic studies was purchased via an NIH shared instrument grant S10 OD030398 (B.W.).

## Author Contributions

Conceptualization, N.M., A.C.B., and J.M.B.; methodology, N.M., A.C.B., A.J.H., V.V., S.D., K.M., R.B., W.J.M., X.Y., M.M., A.L.B., M.A., K.K., A.S., C.T., K.T., Y.Y., I.A., R.Z., Y.Q., B.W., A.M.H., M.D., N.S., M.E.W., M.S., F.C., and J.M.B.; investigation, N.M., A.C.B., A.J.H., V.V., S.D., K.M., R.B., W.J.M., X.Y., M.M., A.L.B., M.A., K.K., A.S., C.T., K.T., Y.Y., I.A., R.Z., Y.Q., B.W., A.M.H., M.D., N.S., M.S., and F.C.; validation, N.M., A.C.B., A.J.H., V.V., S.D., K.M., R.B., W.J.M., X.Y., M.M., A.L.B., M.A., K.K., A.S., C.T., K.T., Y.Y., I.A., R.Z., Y.Q., B.W., A.M.H., M.D., N.S., M.E.W., M.S., and F.C.; formal analysis, N.M., A.C.B., A.J.H., V.V., S.D., K.M., R.B., W.J.M., X.Y., M.M., A.L.B., M.A., K.K., A.S., C.T., K.T., Y.Y., I.A., R.Z., Y.Q., B.W., A.M.H., M.D., N.S., M.E.W., M.S., F.C., and J.M.B.; writing – reviewing and editing, N.M., A.C.B., A.J.H., V.V., S.D., K.M., R.B., W.J.M., X.Y., M.M., A.L.B., M.A., K.K., A.S., C.T., K.T., Y.Y., I.A., R.Z., Y.Q., B.W., A.M.H., M.D., N.S., M.E.W., M.S., F.C. and J.M.B.; funding acquisition, J.M.B.; supervision, J.M.B.

## HIGHLIGHTS

- This work uncovers many novel diet-microbe-host interactions in lipid metabolism.
- Dietary PUFA-driven improvements in metabolic disease depends on gut microbes.
- Gut microbes impact levels of pro-inflammatory and pro-resolving lipid mediators.
- This work is a unique data resource to advance nutrition and microbiome research.

## eTOC BLURB

Diet-microbe-host interactions clearly impact human disease, but how gut microbiota shape metaorganismal lipid metabolism is largely unknown. Mouannes, and Burrows, *et al.* now show that the well-known impact of dietary saturated, monounsaturated, and polyunsaturated fats on cardiometabolic health is uniquely shaped by microbe-host lipid metabolic interactions.

## ABBREVIATIONS USED

AA, arachidonic acid; *Acaca*, acetyl-CoA carboxylase α; ACAT, acyl-CoA:cholesterol acyltransferase; αHYA, 10-hydroxy-*cis*-12,*cis*-15-octadecadienoic acid; αHYB, 10-hydroxy-*cis-* 15-octadecenoic acid; αHYC, 10-hydroxy-*trans*-11,*cis*-15-octadecadienoic acid; αHYD, 13-hydroxy-*cis*-9,*cis*-15-octadecadienoic acid; αHYE, 10,13-dihydroxy-*cis*-15-octadecenoic acid; αKetoA, 10-oxo-*cis*-12,*cis-*15-octadecadienoic acid; αKetoB, 10-oxo-*cis-*15-octadecenoic acid; αKetoC, 10-oxo-*trans*-11,*cis*-15-octadecadienoic acid; αKetoD, 13-oxo-*cis*-9, *cis*-15-octadecadienoic acid; ALA, *cis-9,cis-12,cis*-15-octadecatrienoic acid, α-linolenic acid; BOR, borage oil; CALA1, *cis*-9,*trans*-11,*cis*-15-octadecatrienoic acid; CALA2, *trans*-9,*trans*-11,*cis*-15-octadecatrienoic acid; CALA3, *trans*-10,*cis*-12,*cis*-15-octadecatrienoic acid; CGLA1, *cis*-6,*cis*-9,*trans*-11-octadecatrienoic acid; CGLA2, *cis*-6,*trans*-9,*trans*-11-octadecatrienoic acid; CGLA3, *cis*-6,*trans*-10,*cis*-12-octadecatrienoic acid; CCA, canonical correspondence analysis; CD68, cluster of differentiation 68; CE, cholesteryl ester; Cer, ceramide; CLA1, *cis*-9,*trans*-11-octadecadienoic acid; CLA2, *trans*-9,*trans*-11-octadecadienoic acid; CLA3, *trans*-10,*cis*-12-octadecadienoic acid; CLA-DH, linoleate oxidoreductase; CLA-HY, linoleate hydratase; CLNA, conjugated linolenic acid; CMD, cardiometabolic disease; CROT, carnitine O-octanyltransferase; CVD, cardiovascular disease; c6c9, *cis*-6,*cis*-9-octadecadienoic acid; c6t10, *cis*-6,*trans*-10-octadecadienoic acid; c9c15, *cis*-9,*cis*-15-octadecadienoic acid; DAG, diacylglycerol; DGAT, diacylglycerol acyltransferase; DGLA, di-homo-γ-linolenic acid; DHA, docosahexaenoic acid; DHCR7, 7-dehydrocholesterol reductase; DPA, docosapentaenoic acid; ECH, echium oil; EGFR, epidermal growth factor receptor; ENDOD1, endonuclease domain 1; EPA, eicosapentaenoic acid; EPA-d_5_, *cis*-5,*cis*-8,*cis*-11,*cis*-14,*cis*-17-eicosapentaenoic-19,19,20,20,20-d5 acid; *Fasn*, fatty acid synthase; GBA, glucosylceramidase; GC, Gas Chromatography; GC/MS, Gas Chromatography-Mass Spectrometry; GF, germ-free; γHYA, 10-hydroxy-*cis*-6,*cis*-12-octadecadienoic acid; γHYB, 10-hydroxy-*cis-*6-octadecenoic acid; γHYC, 10-hydroxy-*cis*-6,*trans*-11-octadecadienoic acid; γHYD, 13-hydroxy-*cis*-6,*cis*-9-octadecadienoic acid; γHYE, 10,13-dihydroxy-*cis*-6-octadecenoic acid; γKetoA, 10-oxo-*cis*-6-*cis-*12-octadecadienoic acid; γKetoB, 10-oxo-*cis*6-octadecenoic acid; γKetoC, 10-oxo-*cis*-6,*trans*-11-octadecadienoic acid; γKetoD, 13-oxo-*cis*-6,*cis*-9-octadecadienoic acid; GLA, *cis-6,cis-9,cis*-12-octadecatrienoic acid, γ-linolenic acid; GLP-1, glucagon-like peptide 1; gWAT, gonadal white adipose tissue; HILIC, hydrophilic interaction liquid chromatography; HLS, high linoleic safflower oil; HOS, high oleic acid safflower oil; HYA, 10-hydroxy-*cis*-12-octadecenoic acid; HYARA, 10,12-dihydroxyoctadecanoic acid; HYB, 10-hydroxyoctadecanoic acid; HYC, 10-hydroxy-*trans*-11-octadecenoic acid; HYD, 13-hydroxy-*cis*-9-octadecenoic acid; HYE, 10,13-dihydroxyoctadecanoic acid; KEG1, kidney expressed gene 1; KetoA, 10-oxo-*cis*-12-octadecenoic acid; KetoB, 10-oxo-octadecanoic acid; KetoC, 10-oxo-*tran*s-11-octadecenoic acid; KetoD, 13-oxo-*cis*-9-octadecenoic acid; KetoRA, 12-oxo-*cis*-9-octadecenoic acid; LA, *cis-9-cis*-12-octadecadienoic acid, linoleic acid; LC-MS/MS, liquid chromatography tandem mass spectrometry; LPS, lipopolysaccharide; LTB_4_-d_4_, 5*S*,12*R*-dihydroxy-*cis*-6,*trans*-8,*trans*-10,*cis*-14-eicosatetraenoic-6,7,14,15-d_4_ acid; LXA4, lipoxin A4; MAG, monoacylglycerol; MAMP, microbe-associated molecular pattern; MCP-1, monocyte chemoattractant protein 1; MUFA, monounsaturated fatty acid; MUP1, major urinary protein 1; *N*-DGLA-EA, *N*-dihomo-γ-linolenic acid ethanolamide; NMR, Nuclear Magnetic Resonance; NSAIDs, non-steroidal anti-inflammatory drugs; OA, *cis*-9-octadecenoic acid, Oleic acid; PC, phosphatidylcholine; PDX, protectin DX; PE, phosphatidylethanolamine; PLIN2, perilipin 2; PGE2, prostaglandin E2; PPAR, peroxisome proliferator activated receptor; PRCP, prolylcarboxypeptidase; PUFA, polyunsaturated fatty acid; PYY, peptide YY; RA, 12-hydroxy-*cis*-9-octadecenoic acid; RvD5, resolvin D5; SAMD4A, sterile alpha motif domain-containing protein 4a; SCD-1, stearoyl-CoA desaturase 1; SFA, saturated fatty acid; SL, sphingolipid; SPF, specific pathogen-free; *Srebf1*, sterol regulatory binding element protein 1c; TAG, triacylglycerol; TGFβ, transforming growth factor β; TLR, toll-like receptor; TMEM14C, transmembrane protein 14C; t10c15, *trans*-10,*cis*-15-octadecadienoic acid; UROD, uroporphyrinogen decarboxylase; 5-HEPE, 5-Hydroxyeicosapentaenoic acid; 8,9-DiHETE, 8,9-dihydroxy-eicosatetraenoic acid; 8,9-DiHETrE, 8,9-Dihydroxy-eicosatrienoic acid; 8,15-DiHETE, 8,15-dihydroxyeicosatetraenoic acid; 9-HOTE, 9-hydroxyoctadecatrienoic acid; 10-HODE, 10-Hydroxyoctadecadienoic acid; 10t, *trans*-10-octadecenoic acid; 11,12-DiHETrE, 11,12-dihydroxyeicosatrienoic acid; 12-HODE, 12-hydroxyoctadecadienoic acid; 12-OH, 12-hydroxyoctadecanoic acid; 12-OH-SA, 12-hydroxystearic acid; 12,13-DiHOME,13-dihydroxy-9Z-octadecenoic acid; 12-oxo, 12-oxooctadecanoic acid; 13-KODE, 13-oxooctadeca-9,11-dienoic acid; 14,15-DiHETE, 14,15-dihydroxyeicosatetraenoic acid; 15-deoxy-PGJ2, 15-deoxy-prostaglandin J2; 15(*S*)-HETE-d_8_, 15*S*-hydroxy-*cis*-5,*cis*-8,*cis*-11,*trans*-13-eicosatetraenoic-5,6,8,9,11,12,14,15-d_8_ acid; 16,17-DiDOPE, 16,17-dihydroxy-docosapentaenoic acid; 18S-RvE2, 18S-resolvin E2 (18S-RvE2); 20-HETE, 20-hydroxyeicosatetraenoic acid.

## STAR★Methods

### Key resources table

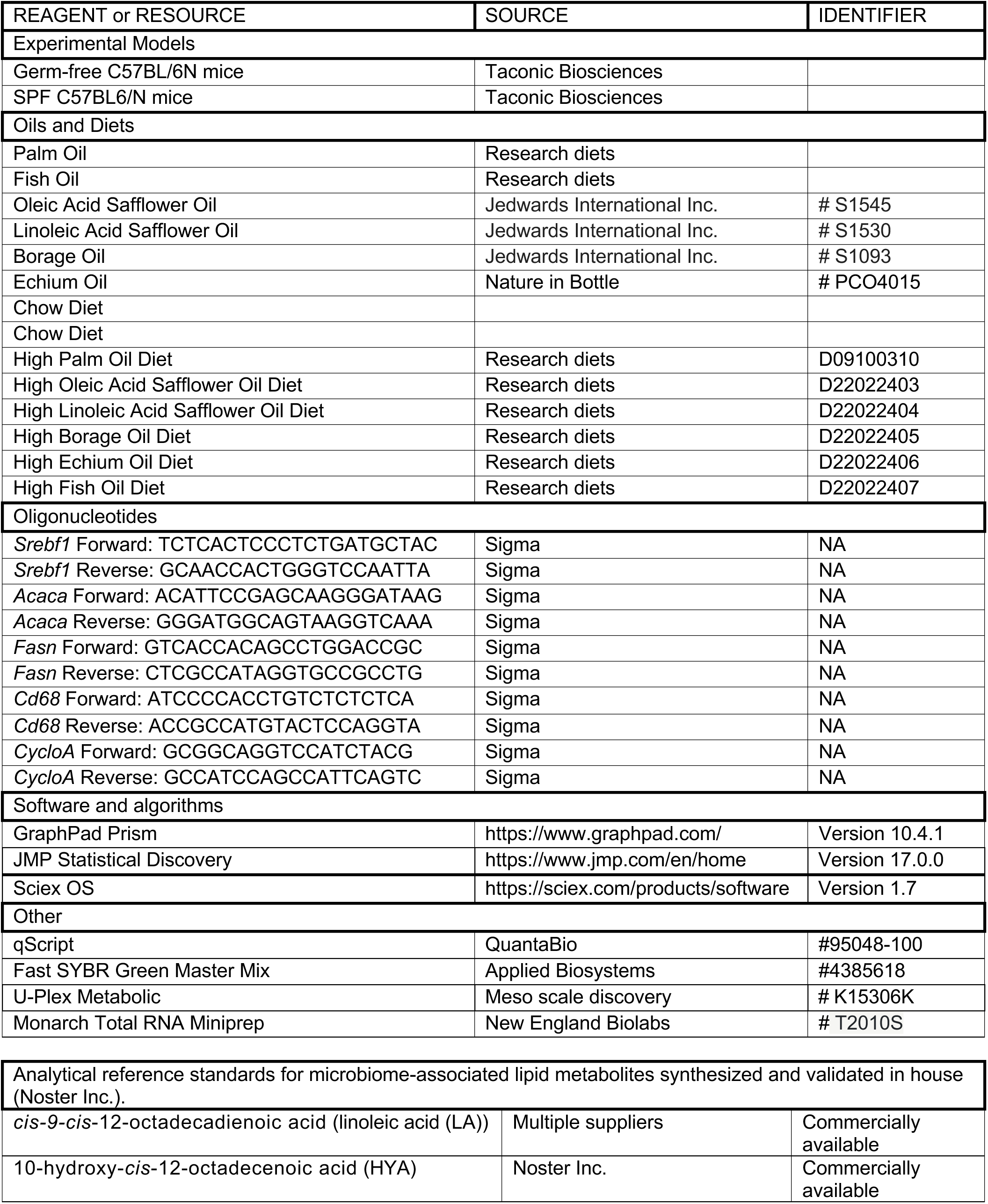
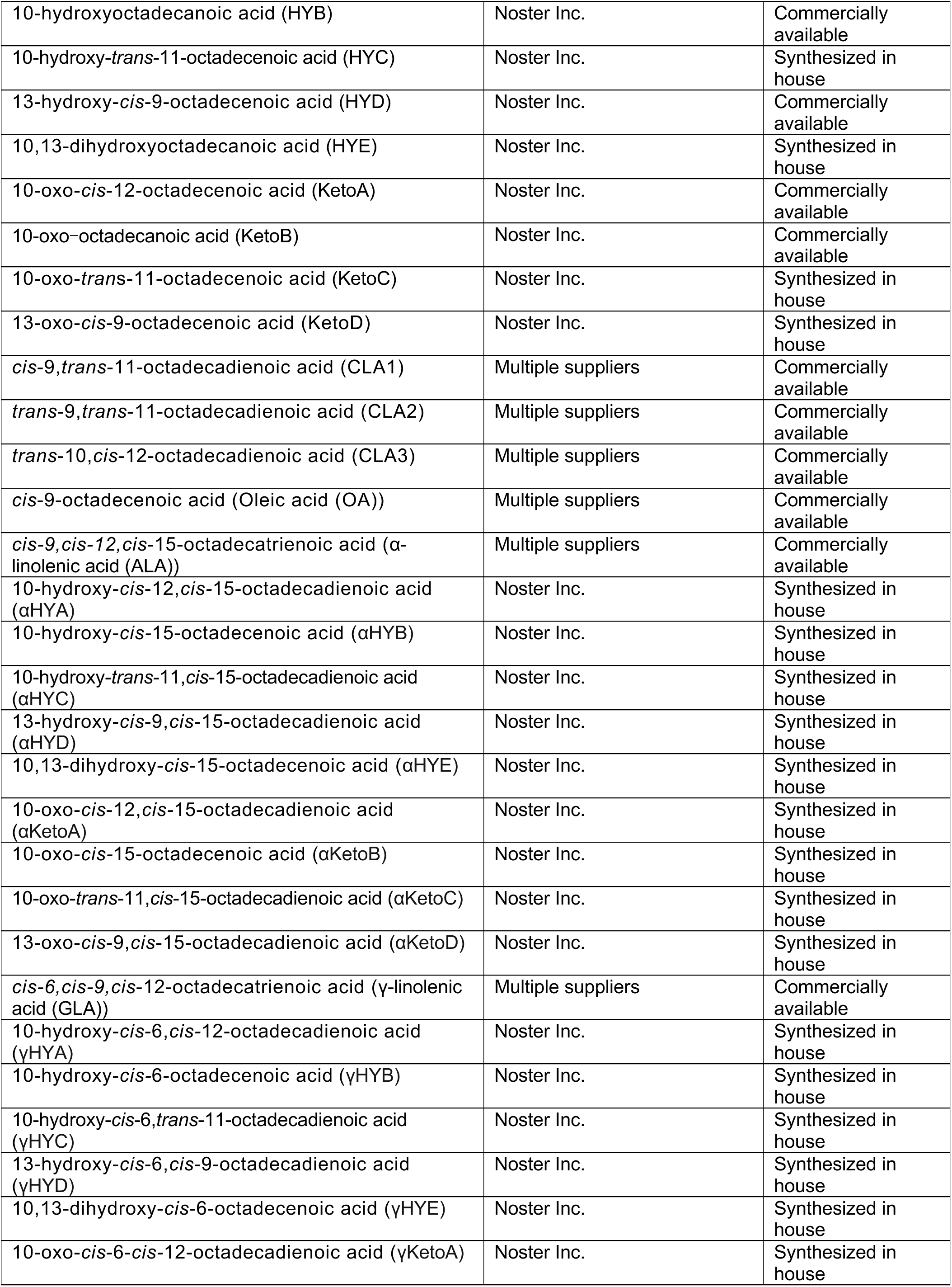
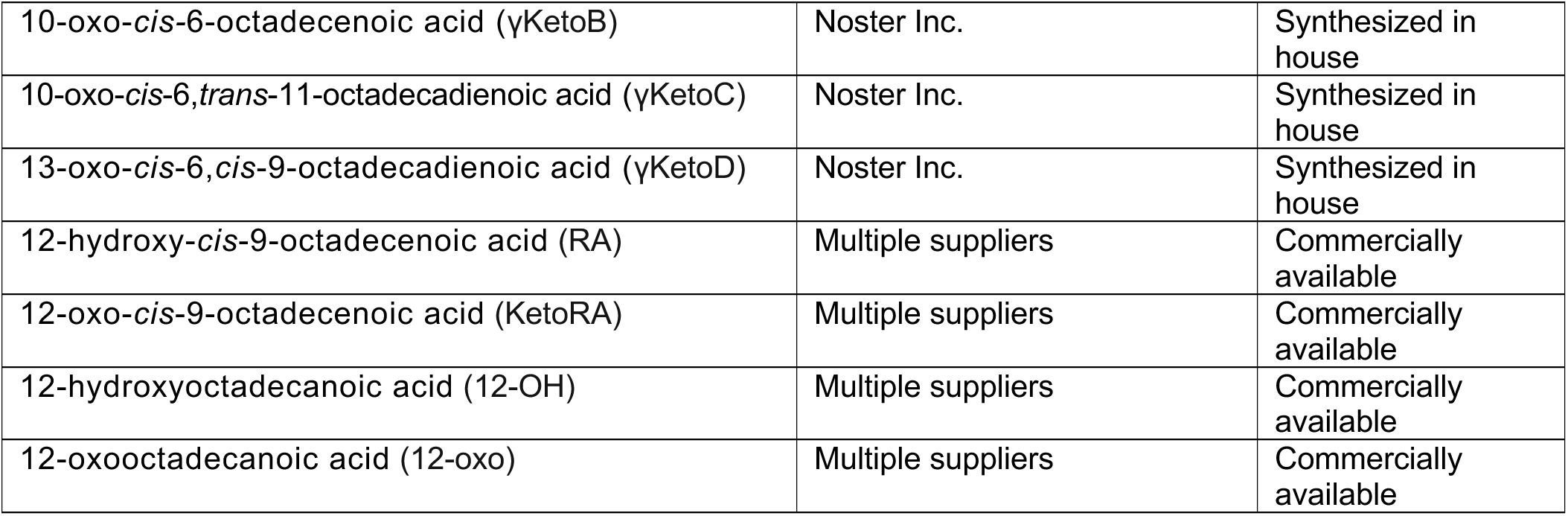

### Animal studies (Mice)

All experiments and procedures were approved by the Cleveland Clinic Lerner Research Institute Institutional Animal Care and Use Committee (IACUC) guidelines protocol # 00002867. Male C57BL6/N mice, either maintained under specific pathogen-free (SPF) or germ-free (GF) conditions, were purchased from Taconic Biosciences at 4 weeks of age and allowed to equilibrate into the Cleveland Clinic Biological Resource Unit (BRU) for an additional 2 weeks before experimental intervention. SPF mice were housed under standard SPF housing conditions within the Cleveland Clinic BRU. To maintain sterility GF mice were housed in hermetically sealed Allentown cages in the Cleveland Clinic Gnotobiotic Facility throughout all dietary interventions. At 6 weeks of age, the mice were switched from sterile chow diet to one of six experimental diets with well-defined fatty acid composition (Figure 1A,1B; Supplemental Table 1). Given we wanted to study the impact of dietary fatty acids under GF settings, we also ensured all experimental diets were sterile (i.e. devoid of any live bacteria) by double irradiation, sterile vacuum packaging, and careful sterile introduction to all mouse cages. The control obesogenic diet (20% protein, 40% carbohydrate, and 40% fat) contained added fructose, high cholesterol, and high fat with a base of palm oil (D09100310, Research Diets, Inc.), which is mostly enriched in SFA including palmitate (16:0) and stearate (18:0). To provide high MUFA substrate, we created a calorically balanced diet that contained high oleic acid (18:1, ω-9) safflower oil (HOS) in the place of the palm oil base in the control diet (D22022403, Research Diets, Inc.). Next, to provide both plant and animal sources of defined ω-6 PUFA substrates we replaced the palm oil base with either high linoleic acid (LA, 18:2, ω-6) safflower oil (HLS; D22022404, Research Diets, Inc.), or GLA (18:3, ω-6)-enriched borage oil (BOR; D22022405, Research Diets, Inc.). Also, to provide plant and animal sources of defined ω-3 PUFA substrates we created matched diets with either ALA (18:3, ω-3)-enriched echium oil (ECH; D22022406, Research Diets, Inc.), or fish oil which has high levels of both EPA (20:5, ω-3) and DHA (22:6, ω-3) (D22022407, Research Diets, Inc). To maintain environmental consistency, both SPF and GF mice were maintained on the same bedding and sterile water throughout the study.

Because special diets that contain casein as a protein source yield false positive 16S rRNA signals due to the presence of killed *Lactococcus*, among others^38^, we first confirmed that our irradiated special diets were indeed sterile. We fed control separate germ-free mice each special diet for 1 week and then switched back to autoclaved chow for 4 weeks. We collected fecal samples before the special diet (Pre), after 1 week of feeding the special diet (On), and then at 2 and 4 weeks post switch back to autoclaved chow (Figure 2A) shows 16S rRNA PCR results from DNA extracted from the fecal samples following methods described by Packey and colleagues^60^. Results from this analysis shown in Figure 2A demonstrates that the 16S rRNA PCR signal that is present when the mice are eating the special diet is not from contamination with live bacteria. Irradiation is less destructive to DNA compared to autoclaving and we also observed Gram + organisms on Gram stains of fecal smears when the mice were on the special diets. These fecal samples were also cultured aerobically on Blood Agar plates (TSA + 5% sheep blood) as well as anaerobically on Brucella agar with hemin and vitamin K plates. No growth was observed on any plates, nor in thioglycollate broth that were also inoculated with fecal slurry while the mice were on the diet as well as 4 weeks after the mice were switched to autoclaved chow. Together, using multiple methods, results demonstrate that the irradiated special diets were sterile.

To maintain colonization status all necropsies were performed under a sterile field contained in a laminar flow biosafety cabinet, with sterile instruments. At necropsy, mice were terminally anesthetized with ketamine/xylazine (100–160 mg/kg ketamine-20–32 mg/kg xylazine), and a midline laparotomy was performed. Blood was collected by cardiac puncture and was immediately processed for plasma isolation by centrifuging at 5500 g, 4°C for 5 mins. The translucent upper layer of plasma was transferred to a fresh cryovial and stored at −80°C. Following blood collection, a whole-body perfusion was conducted by puncturing the inferior vena cava and slowly delivering 10 ml of saline into the heart to remove blood from tissues. Tissues collected included: duodenum, jejunum, ileum, cecum, colon, liver, gonadal white adipose tissue, subscapular brown adipose tissue, kidneys, spleen, and brain. Most tissues were collected and immediately snap frozen in liquid nitrogen for subsequent biochemical analysis or in some cases formalin fixed for histological analysis. It is important to note that although we performed biochemical workup of tissues relevant to cardiometabolic disease (small and large intestine, liver, and adipose tissue), many tissue resources remain and are available to those interested in answering research questions beyond the scope of the current study (please contact the corresponding author to inquire on availability)

### RNA and Realtime PCR Methods

RNA isolation and quantitative polymerase chain reaction (qPCR) was conducted using methods previously described with minor modifications. Liver, gWAT and colon RNA were extracted using the Monarch Total RNA MiniPrep Kit (New England BioLabs, Inc., Cat# T2010S) and cDNA was synthesized using Quanta Bio qScript cDNA Supermix (Cat# 95048-100) in a c1000 Touch Thermal Cycler (Biorad, Hercules, CA). 700ng of cDNA was utilized for Realtime PCR on a 384-well plate, using Applied Biosystem’s Fast SYBR™ Green Master Mix in a Roche LightCycler 480. The levels of induced mRNAs were standardized by the ΔΔCT method and were normalized to cyclophilin A. The specificity of the primers was verified by analyzing the melting curves of the PCR products. Primer information is available in the STAR Methods table.

### Plasma Hormone and Cytokine Quantification

Plasma hormones and cytokine levels were quantified using U-PLEX (product # K15297K) and V-PLEX (K15297K) assays per the manufacturer’s instructions (Meso Scale Diagnostics, Rockville, Maryland, USA).

### Global Proteomics of Mouse Liver

Mouse liver tissue samples were suspended in 150 μL 8M urea Tris-HCl pH8 lysis buffer with freshly added protease inhibitor cocktail. Samples were homogenized by ultrasonication 10s x 3 with 10s between runs. Homogenized samples were centrifuged at 15,000 g for 15 minutes, and the supernatants were transferred to new 1.5 mL tubes. Protein concentrations of the samples were determined by BCA. A 20 μg protein aliquot from each of the samples were reduced by dithiothreitol, alkylated by iodoacetamide and precipitated by cold acetone (−20°C) overnight. Samples were centrifuged at 12,000 g for 15 minutes at 0 °C, and the supernatants were removed. Protein pellets were air dried for 30 minutes and dissolved in 40 μl 50mM tri-ethyl ammonium bicarbonate (TEAB) with 0.3 μg sequencing grade trypsin and incubated at room temperature overnight. Five μg of peptides were taken from each sample and dried in a SpeedVac and reconstituted in 25 µl 0.1% formic acid. Ten µl sample was mixed with 10 µl 0.5x concentration iRT standards and the samples are ready for LCMS analysis. The tissue samples were analyzed using a Data Independent Acquisition (DIA) strategy. For these experiments, a NanoElute HPLC was coupled to a TimsTof Pro2 Q-Tof mass spectrometry system (Bruker Daltonik GmbH, Bremen). The HPLC was equipped with a Bruker 15 cm x 75 µm id C18 ReproSil AQ, 1.9 μm, 120 Å reversed-phase capillary chromatography column. Peptide separation was performed using a 50 minute gradient from 2 to 35% mobile phase B where mobile phase A was 0.1% formic acid and mobile phase B was acetonitrile/0.1% formic acid at a flow rate of 0.3 μL/min. To build a spectral library, 10 μg from each of the digested samples were pooled. The pooled sample was desalted using a Waters C18 Sep-Pak cartridge. The desalted pooled sample was off-line fractionated into 16 fractions using a high pH reversed phase HPLC method on a Waters XBridge BEH C18 2.1 mm X 150 mm, 3.5 μm, 130Å chromatographic column using a 48-minute from 2 to 90% mobile phase B where mobile phase A was 0.1% formic acid 10 mM ammonium formate in water and mobile phase B was 90% acetonitrile/0.1% formic acid 10 mM ammonium formate in water at a flow rate of 0.25 mL/min. Fractions were dried in a SpeedVac and reconstituted in 50 μl 0.1% formic acid each. 10 μl from each fraction was mixed with 10 μl 0.5x concentration iRT standards and the samples were ready for LCMS analysis. For spectral library generation, the fractionated samples were analyzed using a Parallel Accumulation–Serial Fragmentation (PASEF) data dependent acquisition (DDA) method. This method utilized MS1 scans for the identification of precursor ions followed by 10 PASEF MS/MS scans. The tims-MS survey scan was acquired between 0.60 and 1.43 Vs/cm2 and 100–1,700 m/z with a ramp time of 100 ms. The total cycle time for the PASEF scans was 1.17 seconds and the MS/MS experiments were performed with collision energies between 20 eV (0.6 Vs/cm2) and 59 eV (1.6 Vs/cm2). Precursors with 2–5 charges were selected with the target value set to 20,000 a.u. and intensity threshold to 2,500 a.u. Precursors were dynamically excluded for 0.4 minutes.

For protein quantitation, each sample was analyzed using a PASEF data independent acquisition (DIA) method which includes a MS1 full scan followed by MS/MS scans of 32 fixed mass windows between 400 m/z and 1201 m/z. The TIMS-MS survey scan was acquired between 0.60 and 1.43 Vs/cm2 and 100–1,700 m/z with a ramp time of 100 ms. The total cycle time for the PASEF scans was 1.8 seconds and the MS/MS experiments were performed with collision energies between 20 eV (0.85 Vs/cm2) and 59 eV (1.3 Vs/cm2). The spectral library was generated using Pulsar integrated in Spectronaut V17 to search the DDA LC-MS data against the mouse SwissProt protein sequence database (Downloaded on 6-14-2021 with 17050 Entries). Oxidation of Met and protein N-terminal Acetylation were set as variable modifications and carbamidomethylation of Cys was set as fixed modification. The false discovery rate (FDR) of protein, peptide, and peptide spectral match (PSM) were all set to 1% of the identifications from a decoy database generated by reversing the amino acid sequences of the mouse SwissProt protein database. The iRT Reference Strategy was set to Deep learning assisted iRT regression. The DIA data of the samples were searched using Spectronaut V17 software against the spectral library and the mouse SwissProt protein sequence database. The relative abundance of the proteins in these samples was determined using a label free quantitation (LFQ) approach. Protein quantities were expressed as normalized intensities by Spectronaut V17. These are based on the sum of the (raw) intensities of the MS/MS product ions of their peptide precursor peaks that are normalized on total peptide quantities of each sample to ensure that profiles of the abundances across samples accurately reflect the relative amounts of the proteins.

### 16S rRNA Gene Amplicon Sequencing and Dual RNA Sequencing to Examine Gut Microbiota Abundance and Bacterial Gene Expression

Mouse cecum or colon tissue was used to extract the total community RNA using RNeasy Plus Mini Kit (Cat no. 74134). We further enriched a subset of total community RNA for microbial RNA-based sequencing libraries using the MicrobeEnrich Kit (Catalog number AM1901). We further prepared microbial and host RNA sequencing libraries using the QIAseq FX DNA Library Kit (QIAGEN) following our published protocol^61^. We used QIAseq Unique Dual Index Adapters (QIAGEN) for multiplexing. We cleaned ligated products with QIAseq FX Beads (0.8× ratio). We enriched libraries by PCR in 50 µL reactions containing 25 µL of QIAseq HiFi Master Mix, 5 µL of QIAseq IL-Index Primers, and 15 µL of ligated DNA. Cycling conditions were 95 °C for 2 min, followed by 8–12 cycles of 98 °C for 20 s and 65 °C for 30 s, with a final extension at 72 °C for 1 min. We performed size selection using dual QIAseq FX Bead cleanup (0.6× followed by 1.0×) to retain fragments of 300–700 bp. We quantified libraries using the Qubit dsDNA HS Assay and assessed size distribution on a Bioanalyzer High Sensitivity DNA chip. We pooled indexed libraries at equimolar concentrations and performed a final QIAseq FX Bead cleanup (0.8×). We validated the final pool by qPCR using the QIAseq Library Quant Assay Kit and diluted it to 1.8 nM. We denatured the pool with 0.2 N NaOH, neutralized it, and further diluted it to 300 pM. We loaded 1.3 mL onto an Illumina NovaSeq 6000 S4 flow cell and sequenced it using an S4 Reagent Kit (300 cycles) for 2 × 150 bp paired-end sequencing, targeting ∼15 million read pairs per sample. We assessed raw FASTQ quality using FastQC v0.11.9. We trimmed adapters and low-quality bases (Phred < 20) using Trimmomatic v0.40^62^ with parameters: ILLUMINACLIP:QIAseq_adapters.fa:2:30:10, LEADING:20, TRAILING:20, SLIDINGWINDOW:4:20, MINLEN:50. We removed contaminant host reads by mapping against the reference genome (GRCm39) using Bowtie2 v2.4.4 with--very-sensitive-local settings and discarding aligned reads^63^. We performed taxonomic classification of cleaned reads using Kraken2 v2.1.2^64^ with a custom database comprising RefSeq bacterial, archaeal, viral, and fungal genomes. We set the confidence threshold to 0.1. We generated per-sample taxonomic abundance reports using Bracken v2.7^65^ with a read length of 150 bp and k-mer distribution recalculated for our database. We summarized results at genus and species levels and converted raw counts to relative abundances.

We also amplified the V3-V4 hypervariable region of the 16S rRNA gene using methods extensively published by our group earlier^66–71^. Polymerase chain reaction (PCR) was performed in 25 µL reactions containing 12.5 µL of QIAseq 2× HiFi Master Mix (QIAGEN), 0.4 µM of each primer, and 10–50 ng of template DNA. Thermal cycling consisted of initial denaturation at 95 °C for 5 min, followed by 28 cycles of 95 °C for 30 s, 55 °C for 30 s, and 72 °C for 30 s, with a final extension at 72 °C for 5 min. We purified PCR amplicons using QIAquick PCR Purification Kit (QIAGEN) according to the manufacturer’s protocol to remove primers and primer-dimers. We then performed indexing PCR in 25 µL reactions using 5 µL of purified amplicon, 2.5 µL each of QIAseq IL-Forward and IL-Reverse Index Primers (QIAGEN), and 12.5 µL of QIAseq 2× HiFi Master Mix. Cycling conditions were 95 °C for 5 min, followed by 10 cycles of 95 °C for 30 s, 55 °C for 30 s, and 72 °C for 30 s, with a final extension at 72 °C for 5 min. We cleaned indexed libraries using the QIAseq Bead Cleanup Protocol (1.0× ratio) with QIAseq Magnetic Beads (QIAGEN). We quantified individual libraries using the Qubit dsDNA HS Assay Kit (Thermo Fisher Scientific) and assessed fragment size distribution on a Bioanalyzer 2100 using High Sensitivity DNA chips (Agilent Technologies). We pooled equimolar amounts of indexed libraries and performed a final cleanup using QIAseq Beads (0.8× ratio) to remove residual short fragments. We performed quality control (QC) on the pooled library using the QIAseq Library Quant Assay Kit (QIAGEN) on a qPCR instrument to determine exact molarity. We then denatured and diluted the library to 12 pM, spiked with 10 % PhiX control v3 (Illumina), and loaded it onto an Illumina MiSeq-i100 instrument. We demultiplexed raw sequencing data using bcl2fastq v2.20 (Illumina) with default settings. First, we trimmed adapters and low-quality bases (Phred score < 20) using Trimmomatic v0.40^72^. We removed chimeric sequences using the DADA2 consensus method^72^. We assigned taxonomy using the SILVA database (v138.1) with a minimum bootstrap confidence of 80 %. Downstream analysis of 16S rRNA amplicon data was performed using the phyloseq package in R^73^.

Raw reads from host bulk-RNASeq data were quality-trimmed and adapter-removed using Trimmomatic (version 0.39) with default parameters for paired-end reads. Trimmed reads were then aligned to the mouse reference genome (GRCm39) using the STAR aligner (version 2.7.10a^74^) in two-pass mode to generate BAM files. Gene abundance matrices were created by quantifying aligned reads against the Ensembl gene annotation using featureCounts from the Subread package (version 2.0.3)^75^. Differential expression analysis was performed using DESeq2 (version 1.34.0) in R^76^, followed by gene set enrichment analysis (GSEA) for pathway enrichment using the fgsea package (version 1.20.0)^77^ with the KEGG database as the reference, applying a significance threshold of adjusted p-value < 0.05. For multi-omics integration, normalized abundance matrices from host RNA-seq and complementary omics datasets (e.g., proteomics, metatranscriptomics) were analyzed using the DIABLO framework from the mixOmics package (version 6.20.0)^78^, including sparse partial least squares discriminant analysis (sPLS-DA) for feature selection, block integration to identify correlated components across datasets, and generation of circos plots to visualize multi-block correlations and variable contributions.

### Untargeted Metabolomics Analysis

Liver samples were prepared for untargeted metabolomics. Tissues were initially thawed on ice. About 50mg were cut and placed into 1.5mL MPbio lysing matrix D tubes. 600 µL of chilled methanol: water: chloroform (7:2:1 v/v/v) containing 50 µM heavy-labeled internal standards were pipetted into each tube and homogenized using 6×3mm Zirconium beads 3x at high speed (4000 rpm) in 30 second intervals with rest on ice. Blank tubes containing beads with no tissue were also set up as quality check and treated similarly as samples. Both blank and tissue tubes were homogenized. Homogenates were then vortexed for 10 seconds and kept on ice for 5 minutes followed by centrifugation at 10,000 g for 15 min at 4°C. The supernatants were dried overnight in a speed vacuum and dried extracts were re-suspended in water:acetonitrile (95:5 v/v) and normalized based on dilution to control for different tissue weights. Each sample was then randomized and subjected to LC-MS analysis. 2 µL from each sample was taken and pooled and this pooled QC sample was analyzed every 5th injection.

Both Reverse phase and Hydrophilic interaction (HILIC) chromatographic methods were used for untargeted analysis. Reverse phase chromatography was performed by injecting 6 µL of each sample on a Thermo Accucore Vanquish C18 column (100 × 2.1 mm, 1.5 µm) at 60°C coupled to a Thermo Vanquish UHPLC by gradient elution where mobile phase A is 0.1% formic acid in water and mobile phase B is 0.1% formic acid in acetonitrile, flow rate was 0.3 mL/min. The separation was conducted using the following gradient: 0.2 min 5% B; 0.2-1 min, 15% B; 1-8 min, 95% B; 8-10 min, 95% B; 10-12 min, 5% B; 12-15 min, 5% B. HILIC was performed by injecting 6 µL of each sample on a Waters BEH Amide column (2.1×150mm, 2.5 µm) at 40°C coupled to a Thermo Vanquish UHPLC by gradient elution where mobile phase A is 10 mM ammonium acetate with 0.125% acetic acid in water and mobile phase B is 10 mM ammonium acetate with 0.125% acetic acid in 95% acetonitrile. The flow rate was 0.2 mL/min. The separation was conducted using the following gradient: 0 min, 100% B; 0-2 min, 70% B; 2-5.5 min, 60% B; 5.5-11.5 min, 50% B; 11.5-12 min, 100% B; 12-16 min, 100% B. The Orbitrap Q Exactive HF was operated in both positive and negative electrospray ionization modes in different LC-MS runs over a mass range of 56-850 Da using full MS scan at 120,000 resolution. Data dependent acquisitions (DDA) on the pooled representative QC samples include full MS scans at a resolution of 120,000 and HCD MS/MS scans taken on the top 5 most abundant ions at a resolution of 30,000 with dynamic exclusion of 40.0 seconds and the apex trigger set at 2.0 to 4.0 seconds. The resolution of the MS2 scans were taken at a stepped NCE energy of 10.0, 20.0 and 30.0. Data were processed using MS-DIAL (v.4.92) for feature detection, identification and alignment using parameters optimized for data acquired on an Orbitrap mass spectrometer. Metabolites were annotated using spectral libraries from MassBank of North America (MoNA) and MS-DIAL Metabolomics MSP Spectral Kit.

### Quantification of Total Plasma Fatty Acid Levels

To quantify total levels of circulating fatty acids, alkaline hydrolysis of plasma was performed and liberated fatty acid species were quantified by liquid chromatography online tandem mass spectrometry (LC-MS/MS) as previously described^79^. Chemical standards of palmitic acid, palmitoleic acid, stearic acid, oleic acid, linoleic acid, γ-linolenic acid, α-linolenic acid, stearidonic acid, dihomo-γ-linoleic acid, arachidonic acid, eicosapentaenoic acid and docosahexaenoic acid were purchased from Sigma-Aldrich (Vienna, Austria), and the internal standard (heneicosapentaenoic acid, HPA) was purchased from Avanti Polar Lipids, Inc. (Alabaster, Alabama, USA). A 20 μl aliquot of plasma was mixed with 20 μl of internal standard at the concentration of 20 μg/ml and the lipids were extracted into the hexane layer using the Liquid/Liquid extraction method. The hexane layer was dried under nitrogen flow and the pellet was resuspended with 80 μl 75% methanol. Following centrifugation at 12000 rcf for 10 minutes, 40 μl of supernatant was transferred into a HPLC vial and 5 ul was injected on a C18 column (Gemini, 3 μm, 2 x 150mm, Phenomenex) within a UHPLC system (Vanquish, Thermos Fisher Scientific, Waltham, MA, USA) for the separation of fatty acids species. Mobile phases were A (water containing 0.1% acetic acid and 0.03% ammonium hydroxide) and B (methanol/acetonitrile (50/50, v/v) containing 0.1% acetic acid and 0.03% ammonium hydroxide). The run started with 75% mobile phase B from 0 to 2 min at the flow rate of 0.3 ml/min. Solvent B was then increased linearly to 100% B from 2 to 8 min and held at 100% B from 8 to 18 min. The column was finally re-equilibrated with 75% B for 8 min. The HPLC eluent was directly injected into a triple quadrupole mass spectrometer (Quantiva, Thermo Fisher Scientific, Waltham, Massachusetts) and the fatty acids species were ionized at ESI negative mode. Analytes were quantified using Selected Reaction Monitoring (SRM) and the SRM transitions (m/z) were precursor to precursor ions (m/z) at 255 > 255 for palmitic acid, 253 > 253 for palmitoleic acid, 283 > 283 for stearic acid, 281 > 281 for oleic acid, 279 > 279 for linoleic acid, 277 > 277 for γ-linolenic acid and for α-linolenic acid, 275 > 275 for stearidonic acid, 305 > 305 for dihomo-γ-linoleic acid, 303 > 303 for arachidonic acid, 301 > 301 for eicosapentaenoic acid, 327 > 327 for docosahexaenoic acid, 315 > 315 for the internal standard (HPA). The peak areas of the fatty acids were integrated using the software Xcalibur v.4.1 (Thermo Fisher). The Internal standard calibration curves were used for quantitation of fatty acids species in the samples. Resulting data were analyzed using Masslynx Software.

### Global Lipidomics and Oxylipin Quantification in Mouse Liver

Global hepatic lipidomics and oxylipin data were acquired at the UC Davis West Coast Metabolomics Center using methods we have previously described^80,81^. For complex lipid analyses, reverse phased liquid chromatography-tandem mass spectrometry (RPLC-MS/MS) was used to perform lipidomic analysis and began by adding 100 μL run solvent (9: 1 methanol/toluene (v/v)) to microcentrifuge tubes from the dried upper layer of extraction. Chromatography was performed using an Agilent 1290 UHPLC and mass spectra were collected with an Agilent 6550 QTOF mass spectrometer. An Acquity UPLC CSH C18 (100 mm × 2.1 mm, 1.7 μm particle size) column (Waters, Milford, MA) with an Acquity UPLC CSH C18 (5 mm × 1.2 mm, 1.7 μm particle size) pre-column (Waters, Milford, MA) was used with mobile phase A (6: 4 acetonitrile/water (v/v)) and mobile phase B (9: 1 isopropanol acetonitrile (v/v)). Mobile phases A and B were modified with 10 mM ammonium formate and 0.1% formic acid for positive mode ionization and 10 mM ammonium acetate for negative mode ionization. The LC gradient started at 15% B, increased to 30% B from 0–2 minutes, increased to 48% B from 2–2.5 minutes, increased to 82% B from 2.5–11 minutes, increased to 99% B from 11–11.5 minutes, held at 99% B from 11.5–12 minutes, returned to 15% B from 12–12.1 minutes and held at 15% B from 12.1–15 minutes. The autosampler was held at 4 °C and needle wash was performed before and after sample injections for 10 seconds with isopropanol. Injection volumes were 4 μL for both positive and negative mode ionization analyses. Spectral data was collected with a scan range of 120–1700 m/z. MS/MS fragmentation used data-dependent-acquisition (DDA) and was collected for the top 4 most abundant ions from each MS scan. LC-MS/MS data were processed using open source software MS-DIAL (version 4.24) which performed peak picking, deisotoping, automated peak annotation, alignment and gap filling. Blank subtraction was performed by removing features that had a maximum sample intensity/average blank intensity ratio of less than 5. Adduct and duplicate features were flagged using Mass Spectral Feature List Optimizer (MS-FLO) using methods previously described^82,83^. Data from each of the four analytical platforms (RPLC-MS/MS ESI+/−, HILIC-MS/MS ESI+/−) were processed separately and combined after data curation. No data normalization was performed because no trend in data intensities was observed from the internal standards during data acquisition. Peak height was used for all quantitation. Metabolite annotations were made using defined confidence levels based on accurate mass, MS/MS library matching to experimental data, and retention time from authentic standards run on the same instrument^82,83^. Tandem MS/MS libraries of the MassBank of North America (<u>MassBank.us</u>) and NIST17 (NIST, Gaithersburg, MD) were used for spectral matching. A manual curation of datasets was performed to reduce in-source fragment annotations identified by very similar RT and high correlation between features. Manual review of MS/MS matches was performed to remove poor spectral matches since false positive annotations can occur when automatically matching MS/MS from complex biological samples to large MS/MS spectral libraries^82,83^. For oxylipin analyses, samples were added to a 96-well plate followed sequentially by 25 μL anti-oxidant solution, 25 μL of surrogate standards in methanol, 25 μL of CUDA and PHAU standards in methanol, and 125 μL acetonitrile/methanol 1:1. Plates were then vortexed for 30 s, centrifuged at 6 °C for 5 min at 15,000 rcf and filtered through a PVDC 0.2-micron filter plate. Plates were then sealed and kept at −20 °C until analysis. Extracted oxylipins were separated and quantified using a Waters i-Class Acquity UHPLC system coupled to a Sciex 6500+ QTRAP mass spectrometer operated in negative ionization mode. Oxylipins were quantified by targeted, retention time-specific, multiple reaction monitoring ion transitions. A total of 78 oxylipins were targeted, and targets that appeared in any sample above the signal-to-noise ratio of 3:1 were quantified. This resulted in quantified values for 27 ω-6-related oxylipins, including 17 AA-related and 10 LA-related oxylipins. There were 12 ω-3-related oxylipins measured, which included 6 ALA-related, 4 DHA-related, and 2 EPA-related oxylipins. Oxylipins measured in a sample below the limit of quantitation (LOQ), defined as signal-to-noise ratio below 3:1, were converted to 10% of the LOQ.

### Targeted Quantification of Select Pro-Inflammatory and Pro-Resolving Lipid Mediators in the Liver by Liquid Chromatography Tandem Mass Spectrometry (LC-MS/MS)

Livers were minced in ice-cold methanol containing internal deuterium-labeled standards and stored at −80°C to allow for protein precipitation. The deuterated internal standards included d_5_-RvD2, d_5_-17R-RvD1, d_5_-RvD3, d_5_-MaR1, d_5_-MaR2, d_4_-RvE1, d_4_-LTB_4_, d_8_-5-HETE, d_8_-12-HETE, d_8_-15-HETE, d_5_-17-HDHA, d_4_-PGE_2_ and d_5_-LXA_4,_ (Cayman Chemical) and were used to assess extraction recovery and retention time. The tissue samples were centrifuged (3,000 rpm) and the supernatants were subjected to solid phase extraction and LC-MS/MS analysis. Briefly, lipid mediators were extracted by C18 column chromatography and were eluted in methyl formate fractions as described^82^. The solvent was evaporated under a gentle stream of N_2_ gas and lipid mediators were resuspended in methanol:water (50:50). For analysis, a high performance liquid chromatograph (HPLC; Shimadzu) equipped with a reverse-phase (C18) column (100 x 4.6mm, 2.6μm; held at 50°C using a column oven) was operated using a mobile phase consisting of LC-MS grade methanol:water:acetic acid (50:50:0.01 v/v/v) that was ramped to 98:2:0.01 (v/v/v) at a constant flow rate of 0.5mL/min. The HPLC was coupled to a QTrap6500+ mass spectrometer (Sciex) operating in negative ionization mode using Analyst software v.1.7 in part as described^83^. Identification and quantification of lipid mediators was achieved using multiple reaction monitoring transitions established for each mediator. The levels of individual lipid mediators were quantified based on calibration curves using external standards for each mediator followed by normalization to extraction recovery of internal deuterium-labeled standards with similar chromatographic retention times using Sciex OS (v1.7). Criteria for quantification included matching of retention time to synthetic standards acquired in parallel, a chromatographic peak that consisted of at least 8-10 data points, a signal:noise ratio > or equal to 5, and above the lower limit of quantification (defined as a coefficient of variation of replicate injections of <20% based on peak areas of synthetic standards). Final levels of lipid mediators were normalized to the tissue weight of each sample.

### Quantification of Gut Microbe-Derived PUFA Metabolites in Plasma, Colon, and Liver

All listed microbiome-associated lipid metabolites were quantified by liquid chromatography tandem mass spectrometry (LC-MS/MS) using analytical reference standards and quantitative protocols developed and validated under the Noster analytical standard protocol (Noster Inc., Kyoto, Japan), as described below^84^. The chemical structures of the synthesized analytical reference standards were identified by Gas Chromatography-Mass Spectrometry (GC/MS) and Nuclear Magnetic Resonance (NMR), and their purity were determined by Gas chromatography prior to use for calibration, and the synthesized analytical standards are maintained under controlled production records to ensure future batch reproducibility. Metabolites were quantified by a Nexera X3 HPLC system (Shimadzu Corporation, Kyoto, Japan) coupled with an LCMS-8060NX triple quadrupole mass spectrometer (Shimadzu Corporation) using electrospray ionization (ESI) in negative ion mode. Data processing and peak integration were performed using Labsolutions INSIGHT software (Shimadzu Corporation) consistent parameters to ensure analytical reproducibility and comparability across samples.

| List of lipids and quantified microbiome-associated lipid metabolites and information for LC-MS/MS analysis<br>(IS: Internal Standard) |  |  |  |  |
| --- | --- | --- | --- | --- |
| Compound Name / Lot No. of Noster inc.<br>analytical reference standard | Retention time<br>(min) | Precursor ion<br>(m/z) | Product ion<br>(m/z) | Collision energy<br>(eV) |
| *cis*-9-*cis*-12-octadecadienoic acid (linoleic acid (LA)) / Commercial product | 29.0 | 279.3 | 261.3 | 18.0 |
| 10-hydroxy-*cis*-12-octadecenoic acid (HYA) / SN15 | 20.9 | 297.4 | 185.2 | 20.0 |
| 10-hydroxyoctadecanoic acid (HYB) / TC03 | 23.5 | 299.3 | 141.3 | 23.0 |
| 10-hydroxy-*trans*-11-octadecenoic acid (HYC) / UN08 | 21.1 | 297.2 | 185.3 | 20.2 |
| 13-hydroxy-*cis*-9-octadecenoic acid (HYD) / TG07 | 20.7 | 297.2 | 197.3 | 20.0 |
| 10,13-dihydroxyoctadecanoic acid (HYE) / SA14 | 12.5 | 315.3<br>315.3 | 185.3<br>197.3 (Qualifier) | 26.0<br>25.0 |
| 10-oxo-*cis*-12-octadecenoic acid (KetoA) / QM07 | 21.6 | 295.4 | 139.3 | 20.0 |
| 10-oxo-octadecanoic acid (KetoB) / TC10 | 24.1 | 297.3 | 139.3 | 21.0 |
| 10-oxo-*trans*-11-octadecenoic acid (KetoC) / UK08 | 22.1 | 295.4 | 139.3 | 22.0 |
| 13-oxo-*cis*-9-octadecenoic acid (KetoD) / RH02 | 21.4 | 295.4 | 179.2 | 20.0 |
| *cis*-9,*trans*-11-octadecadienoic acid (CLA1) / YH01 | 29.6 | 279.4 | 261.4 | 18.0 |
| *trans*-9,*trans*-11-octadecadienoic acid (CLA2) / SK02 | 30.9 | 279.3 | 279.3 | 11.0 |
| *trans*-10,*cis*-12-octadecadienoic acid (CLA3) / YH03 | 29.6 | 279.3 | 261.3 | 19.0 |
| *cis*-9-octadecenoic acid (Oleic acid (OA)) / Commercial product | 34.0 | 281.4 | 281.4 | 25.0 |
| *trans*-10-octadecenoic acid (10t) / SK02 | 34.9 | 281.4 | 281.4 | 20.0 |
| *cis*-9,*cis*-12,*cis*-15-octadecatrienoic acid (α-linolenic acid (ALA)) / Commercial product | 26.1 | 277.3 | 233.3 | 14.0 |
| 10-hydroxy-*cis*-12,*cis*-15-octadecadienoic acid (αHYA) TN27 | 18.7 | 295.3 | 185.3 | 18.0 |
| 10-hydroxy-*cis*-15-octadecenoic acid (αHYB) / RH07 | 20.6 | 297.3 | 141.1 | 23.0 |
| 10-hydroxy-*trans*-11,*cis*-15-octadecadienoic acid (αHYC) / SM20 | 18.7 | 295.4 | 185.3 | 19.0 |
| 13-hydroxy-*cis*-9,*cis*-15-octadecadienoic acid (αHYD) / TH21 | 18.8 | 295.4 | 225.3 | 20.0 |
| 10,13-dihydroxy-*cis*-15-octadecenoic acid (αHYE) / QC16 | 9.1 | 313.2<br>313.2 | 243.3<br>185.3 (Qualifier) | 21.0<br>23.0 |
| 10-oxo-*cis*-12,*cis*-15-octadecadienoic acid (αKetoA) / UA16 | 19.6 | 293.4 | 139.2 | 20.0 |
| 10-oxo-*cis*-15-octadecenoic acid (αKetoB) / WH13 | 21.4 | 295.4 | 155.3 | 21.0 |
| 10-oxo-*trans*-11,*cis*-15-octadecadienoic acid (αKetoC) / WD04 | 19.6 | 293.3 | 139.2 | 19.0 |
| 13-oxo-*cis*-9,*cis*-15-octadecadienoic acid (αKetoD) / SA06 | 19.6 | 293.3 | 205.3 | 19.0 |
| *cis*-9,*trans*-11,*cis*-15-octadecatrienoic acid (CALA1) / XA09 | 26.3 | 277.3 | 233.3 | 17.0 |
| *trans*-9,*trans*-11,*cis*-15-octadecatrienoic acid (CALA2) / RJ15 | 27.1 | 277.2 | 233.35 | 16.0 |
| *trans*-10,*cis*-12,*cis*-15-octadecatrienoic acid (CALA3) / XB21 | 26.5 | 277.3 | 233.3 | 17.0 |
| *cis*-9,*cis*-15-octadecadienoic acid (c9c15) / RL03 | 29.0 | 279.4 | 261.4 | 18.0 |
| *trans*-10,*cis*-15-octadecadienoic acid (t10c15) / RL02 | 29.8 | 279.4 | 279.4 | 20.0 |
| *cis*-6,*cis*-9,*cis*-12-octadecatrienoic acid (γ-linolenic acid (GLA)) / Commercial product | 26.5 | 277.3 | 233.3 | 15.0 |
| 10-hydroxy-*cis*-6,*cis*-12-octadecadienoic acid (γHYA) / PC13 | 19.8 | 295.4 | 183.2 | 19.0 |
| 10-hydroxy-*cis*-6-octadecenoic acid (γHYB) / RJ20 | 22.0 | 297.1 | 155.2 | 20.0 |
| 10-hydroxy-*cis*-6,*trans*-11-octadecadienoic acid (γHYC) / QK30 | 19.8 | 295.4<br>295.4 | 155.3<br>183.3 (Qualifier) | 18.0<br>19.0 |
| 13-hydroxy-*cis*-6,*cis*-9-octadecadienoic acid (γHYD) / XE22 | 19.1 | 295.4 | 195.3 | 18.0 |
| 10,13-dihydroxy-*cis*-6-octadecenoic acid (γHYE) / QL09 | 11.0 | 313.3<br>313.3 | 155.2<br>195.3 (Qualifier) | 23.0<br>23.0 |
| 10-oxo-*cis*-6-*cis*-12-octadecadienoic acid (γKetoA) / YK09 | 20.5 | 293.3 | 137.2 | 20.0 |
| 10-oxo-*cis*-6-octadecenoic acid (γKetoB) / SD02 | 22.6 | 295.4 | 155.3 | 21.0 |
| 10-oxo-*cis*-6,*trans*-11-octadecadienoic acid (γKetoC) / RE03 | 20.8 | 293.3 | 137.2 | 20.0 |
| 13-oxo-*cis*-6,*cis*-9-octadecadienoic acid (γKetoD) / SA06 | 19.8 | 293.3 | 195.3 | 19.0 |
| *cis*-6, *cis*-9, *trans*-11-octadecatrienoic acid (CGLA1) / RL10 | 27.1 | 277.2 | 233.35 | 15.0 |
| *cis*-6, *trans*-9, *trans*-11-octadecatrienoic acid (CGLA2) / RJ15 | 28.0 | 277.3 | 277.3 | 7.0 |
| *cis*-6, *trans*-10, *cis*-12-octadecatrienoic acid (CGLA3) / SM16 | 26.7 | 277.2 | 233.35 | 16.0 |
| *cis*-6,*cis*-9-octadecadienoic acid (c6c9) / RJ15 | 30.0 | 279.4 | 261.4 | 16.0 |
| *cis*-6,*trans*-10-octadecadienoic acid (c6t10) / YL16 | 30.6 | 279.4 | 279.4 | 9.0 |
| 12-hydroxy-*cis*-9-octadecenoic acid (RA) / RN04 | 20.8 | 297.2 | 183.3 | 18.0 |
| 12-oxo-*cis*-9-octadecenoic acid (KetoRA) / RM25 | 21.4 | 295.4 | 165.3 | 20.0 |
| 10,12-dihydroxyoctadecanoic acid (HYARA) / RK15 | 17.1 | 315.2<br>315.2 | 185.2<br>199.2 (Qualifier) | 23.0<br>24.0 |
| 12-hydroxyoctadecanoic acid (12-OH) / RN04 | 22.8 | 299.3 | 169.3 | 23.0 |
| 12-oxooctadecanoic acid (12-oxo) / RN04 | 23.5 | 297.2 | 167.2 | 21.0 |
| IS: 15*S*-hydroxy-*cis*-5,*cis*-8,*cis*-11,*trans*-13-eicosatetraenoic-5,6,8,9,11,12,14,15-$d_8$ acid (15(*S*)-HETE-$d_8$) / Commercial product | 19.9 | 327.4 | 226.3 | 13.0 |
| IS: 5*S*,12*R*-dihydroxy-*cis*-6,*trans*-8,*trans*-10,*cis*-14-eicosatetraenoic-6,7,14,15-$d_4$ acid (LTB<sub>4</sub>-$d_4$) / Commercial product | 13.5 | 339.4 | 197.3 | 15.0 |
| IS: *cis*-5,*cis*-8,*cis*-11,*cis*-14,*cis*-17-eicosapentaenoic-19,19,20,20,20-$d_5$ acid (EPA-$d_5$) / Commercial product | 25.5 | 306.4 | 262.4 | 11.0 |

Samples from plasma, colon, and liver were, minced if needed, stored in methanol at −30 °C overnight. The sample solutions were then centrifuged at 3,000×g, and the supernatants were collected. The supernatants were mixed with deuterium-labeled internal standards (15(*S*)-HETE-d_8_, LTB_4_-d_4_ and EPA-d_5_) for the purpose of monitoring extraction efficiency and instrument performance. The mixed solutions were subjected to solid-phase extraction (Monospin AX-18, GL Sciences Inc., Tokyo, Japan) to obtain the injection solution for LC-MS/MS. Liquid chromatography was performed by injecting 5µL volume of injection solution prepared from plasma sample (in case of injection solutions prepared from colon or liver samples, injection volume was 1µL each) on an Acquity UPLC BEH C18 (1.0×150 mm, 1.7 μm, Waters, Milford, Massachusetts, USA) analytical column at 35°C by gradient elution where mobile phase A is 0.1% acetic acid in water and mobile phase B is acetonitrile/methanol (4:1, v/v). The separation was conducted using the following gradient elution program: 50% B for 5.0 min, 50-70% B from 5.0 to 15.0 min, 70-80% B from 15.0 to 25.0 min, hold 80% B from 25.0 to 33.0 min, 80-100% B from 33.0 to 35.0 min, hold 100% B from 35.0 to 45.0 min, and hold 50% B for 5.0 min for column equilibration with flow rates of 0.05 mL/min (0-25 min), 0.08 mL/min (25-33 min), 0.1 mL/min (33-45 min), and 0.05mL/min (45-50 min).

Absolute quantification was performed using calibration curves generated from serial dilutions of LC-MS/MS reference standards for microbiome-associated lipid metabolites developed under Noster analytical standard protocol analyzed independently of the biological samples. Through the validation of the analytical method using adequate concentration sample solutions of LC-MS/MS reference standards for microbiome-associated lipid metabolites developed under Noster analytical standard protocol, it was confirmed that the calibration curve for each microbiome-associated lipid metabolite showed good linearity within the range of 0.1-1,000 pg/μL (the coefficient of determination (R^2^) were more than 0.99 for all calibration curves). While comprehensive recovery and matrix-effect studies were beyond the scope of this resource-focused study, internal standard–based normalization ensured quantitative consistency across samples. The accuracies and precisions of the analytical method for microbiome-associated lipid metabolites using 3 different concentration solutions prepared and repeated 3 times showed that the accuracies were between 80 and 120%, and the RSDs of precision were less than 20%.

### Leveraging the Untargeted Metabolomics Datasets to Determine Alterations in *N*-Acyl Lipids

The untargeted data generated by Orbitrap C18 positive mode were mined for the relative abundance of diverse *N*-acyl lipids using methods recently described by Mannochio-Russo and colleagues^46^. The raw data files were processed in MS-Dial (version 4.9) for peak detection, deconvolution, alignment and MS/MS spectral extraction. The exported files were subjected to feature based molecular network flow in GNPS2. The precursor and MS/MS fragment ion tolerances were set to 0.02 Da. Spectral library matching was conducted against the GNPS *N*-acyl lipid MassQL library^44^. The parameters for the *N*-acyl lipids library search were set to have a cosine value above 0.7 and at least 6 matched fragments. The molecular networks were visualized in Cytoscape (version 3.10.4)

### Standard Statistical Analyses

All data are expressed as the mean ± standard error of the mean (S.E.M.). Other than 16S, proteomics, and RNA sequencing data which are described above, all data were analyzed using ANOVA with post-hoc Tukey-Kramer HSD for all pairs comparisons. Differences were considered significant at *p*<0.05. All group means comparisons were performed using JMP statistical discovery software (version 17.0.0; SAS Institute).

## ONLINE SUPPLEMENT

Submitted in association with this body of work is an online supplement with 7 additional supplemental figures that are related to the 7 main figures in the main manuscript file.

## ONLINE SUPPLEMENT FILES (All provided as accessory excel data tables)

**1. Supplemental Table 1.** Diet information sheet including ingredient list.
**2. Supplemental Table 2.** Untargeted lipidomics analyses of mouse liver using reverse phase liquid chromatography high resolution tandem mass spectrometry (RPLC-MS/MS).
**3. Supplemental Table 3.** Targeted lipidomic analysis of gut bacterially-produced fatty acid metabolites in the plasma, liver, and distal small intestine (ileum) via liquid chromatography tandem mass spectrometry (LC-MS/MS).
**4. Supplemental Table 4.** Targeted lipidomic analysis of diverse oxylipin species in mouse liver was performed using ultra high performance liquid chromatography tandem mass spectrometry (UHPLC-MS/MS).
**5. Supplemental Table 5.** Quantitative global proteomics in mouse liver was performed using Data Independent Acquisition (DIA) liquid chromatography tandem mass spectrometry (LC-MS/MS).
**6. Supplemental Table 6.** Untargeted metabolomics in mouse liver data using C18 column separation coupled to high resolution mass spectrometry.
**7. Supplemental Table 7.** Untargeted metabolomics in mouse liver data using HILIC column separation coupled to high resolution mass spectrometry.

## ONLINE SUPPLEMENTAL INFORMATION

**Figure S1.**
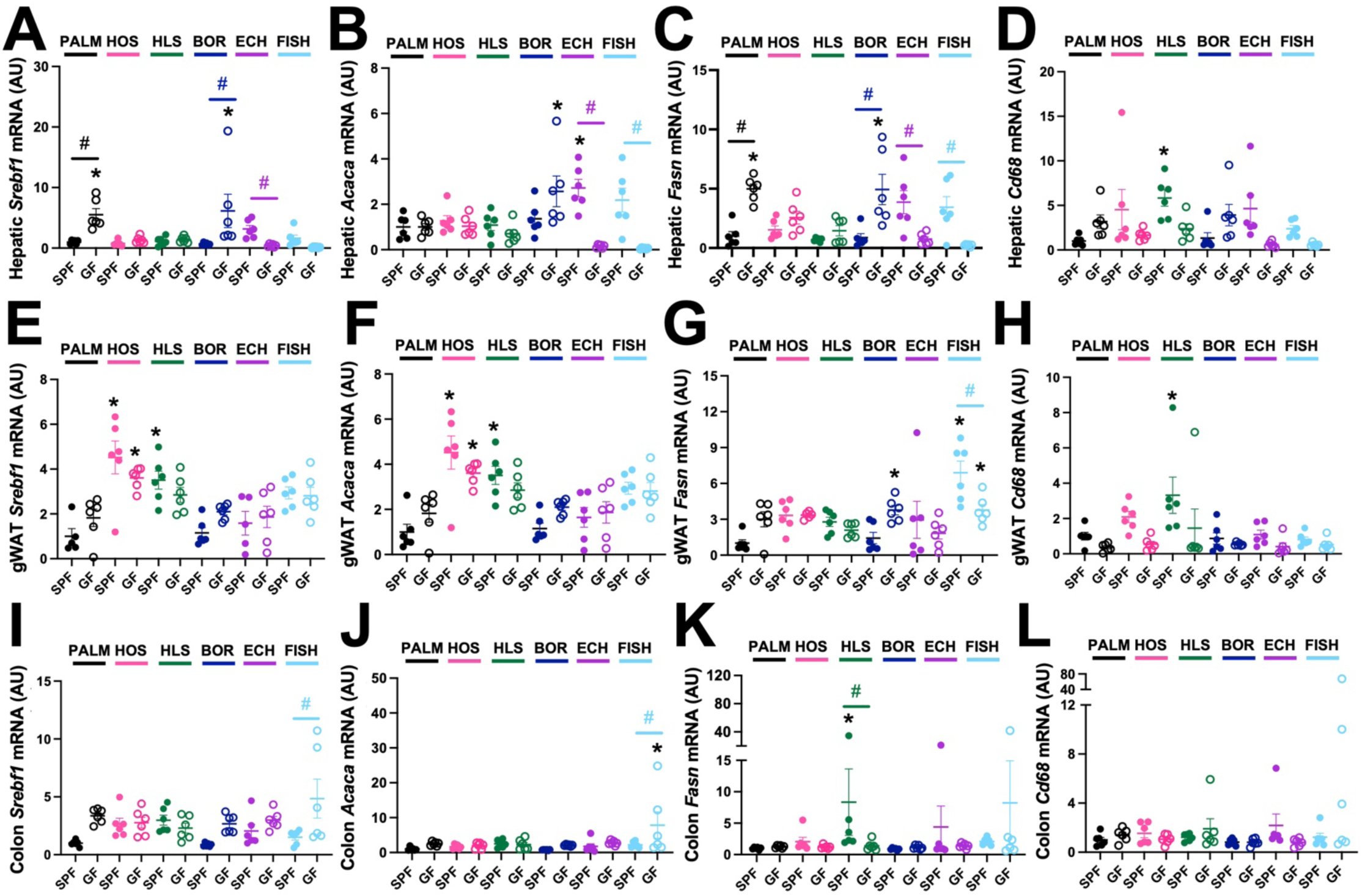
The Ability of Dietary Fatty Acids to Alter Host Metabolic Gene Expression is Shaped by Resident Microbiota (*Related to* Figure 1). Starting at 6 weeks of age, either conventionally-raised specific pathogen-free (SPF) or germ-free (GF) cohorts of male C57BL/6N mice were maintained on sterile fatty acid-defined diets containing saturated (Palm), high oleic safflower (HOS), high linoleic safflower (HLS), borage (BOR), echium (ECH), or fish oils for 18 weeks. Thereafter, **(A-D)** liver, **(E-H)** gonadal white adipose (gWAT), and **(I-L)** colon tissue was subjected to quantitative real-time PCR (qPCR) to examine the mRNA expression of sterol regulatory element-binding protein 1c (*Srebf1*), acetyl-CoA carboxylase α (*Acaca*), fatty acid synthase (*Fasn*), and cluster of differentiation 68 (*Cd68*). Data represent the mean ± S.E.M. from n=6 per group, and statistically significant differences (*p*<0.05) were detected using ANOVA with post-hoc Tukey-Kramer HSD for all pairs comparisons. * = significantly different than the SPF palm oil-fed control group; # = significantly different when comparing SPF and GF groups within each dietary condition.

**Figure S2.**
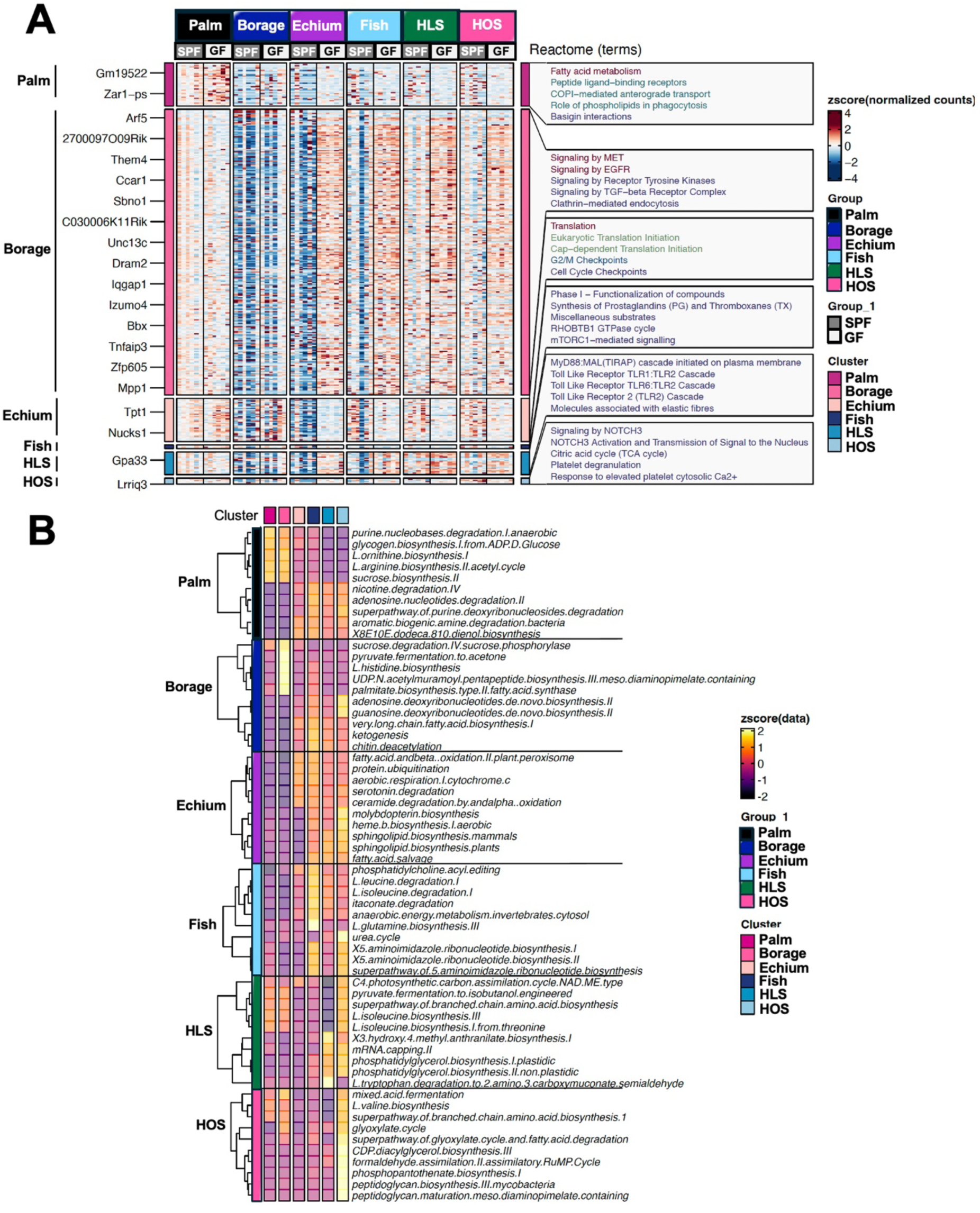
Metatranscriptomic Analyses of Colon Tissue Reveal (*Related to* Figures 1 *and 2*). Starting at 6 weeks of age, either conventionally-raised specific pathogen-free (SPF) or germ-free (GF) cohorts of male C57BL/6N mice were maintained on sterile fatty acid-defined diets containing saturated (Palm), high oleic safflower (HOS), high linoleic safflower (HLS), borage (BOR), echium (ECH), or fish oils for 18 weeks. Thereafter, colon tissue was subject to shotgun metatranscriptomic sequencing with data analytic pipelines to examine both host reads (A) and bacterial reads (B). Data represent the mean ± S.E.M. from n=6 per group, and statistically significant differences (*p*<0.05) were detected using ANOVA with post-hoc Tukey-Kramer HSD for all pairs comparisons. * = significantly different than the SPF palm oil-fed control group; # = significantly different when comparing SPF and GF groups within each dietary condition.

**Figure S3.**
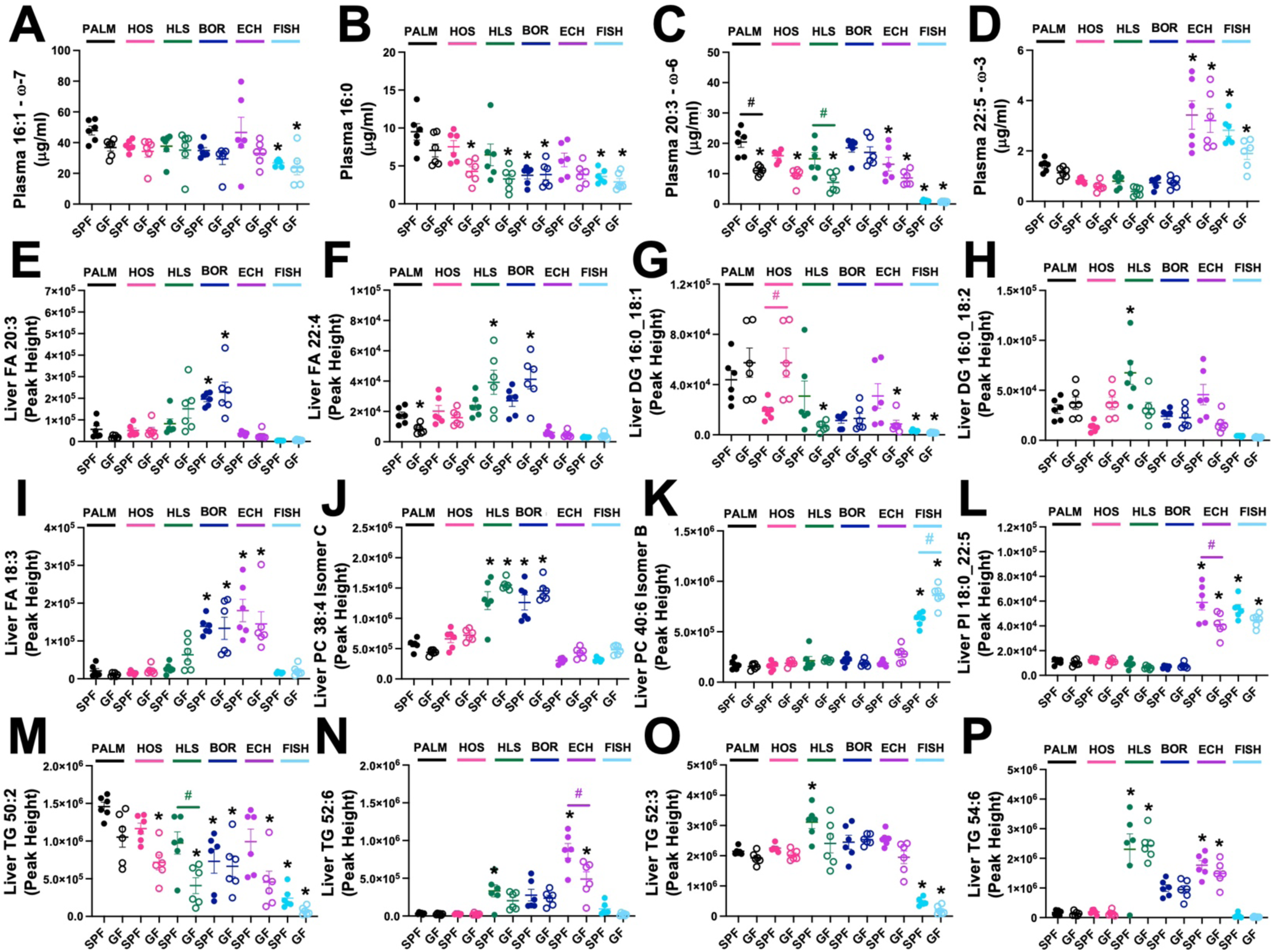
Diet-Microbe-Host Interactions Influence the Systemic Lipidome Within the Mouse Metaorganism (*Related to* Figure 3). Starting at 6 weeks of age, either conventionally-raised specific pathogen-free (SPF) or germ-free (GF) cohorts of male C57BL/6N mice were maintained on sterile fatty acid-defined diets containing saturated (Palm), high oleic safflower (HOS), high linoleic safflower (HLS), borage (BOR), echium (ECH), or fish oils for 18 weeks. Thereafter, plasma and liver samples were subjected to lipidomic analyses to examine the levels of total fatty acids in the circulation as well as diverse complex lipids in the liver. **(A-D)** The total levels of plasma fatty acids including **(A)** palmitoleic acid (16:1, ω-7), **(B)** palmitic acid (16:0) **(C)** di-homo-γ-linolenic acid (20:3, ω-6), and **(D)** docosapentaenoic acid (22:5, ω-3) was quantified using liquid chromatography tandem mass spectrometry (LC-MS/MS). **(E-P)** Hepatic levels of select molecular species of unesterified fatty acids (FA) and esterified lipids including phosphatidylcholines (PC), phosphatidylinositols (PI), diacylglycerols (DAG), and triacylglycerols (TG) were quantified using reverse phase liquid chromatography – high resolution tandem mass spectrometry (RPLC-MS/MS). Data represent the mean ± S.E.M. from n=6 per group, and statistically significant differences (*p*<0.05) were detected using ANOVA with post-hoc Tukey-Kramer HSD for all pairs comparisons. * = significantly different than the SPF palm oil-fed control group; # = significantly different when comparing SPF and GF groups within each dietary condition.

**Figure S4.**
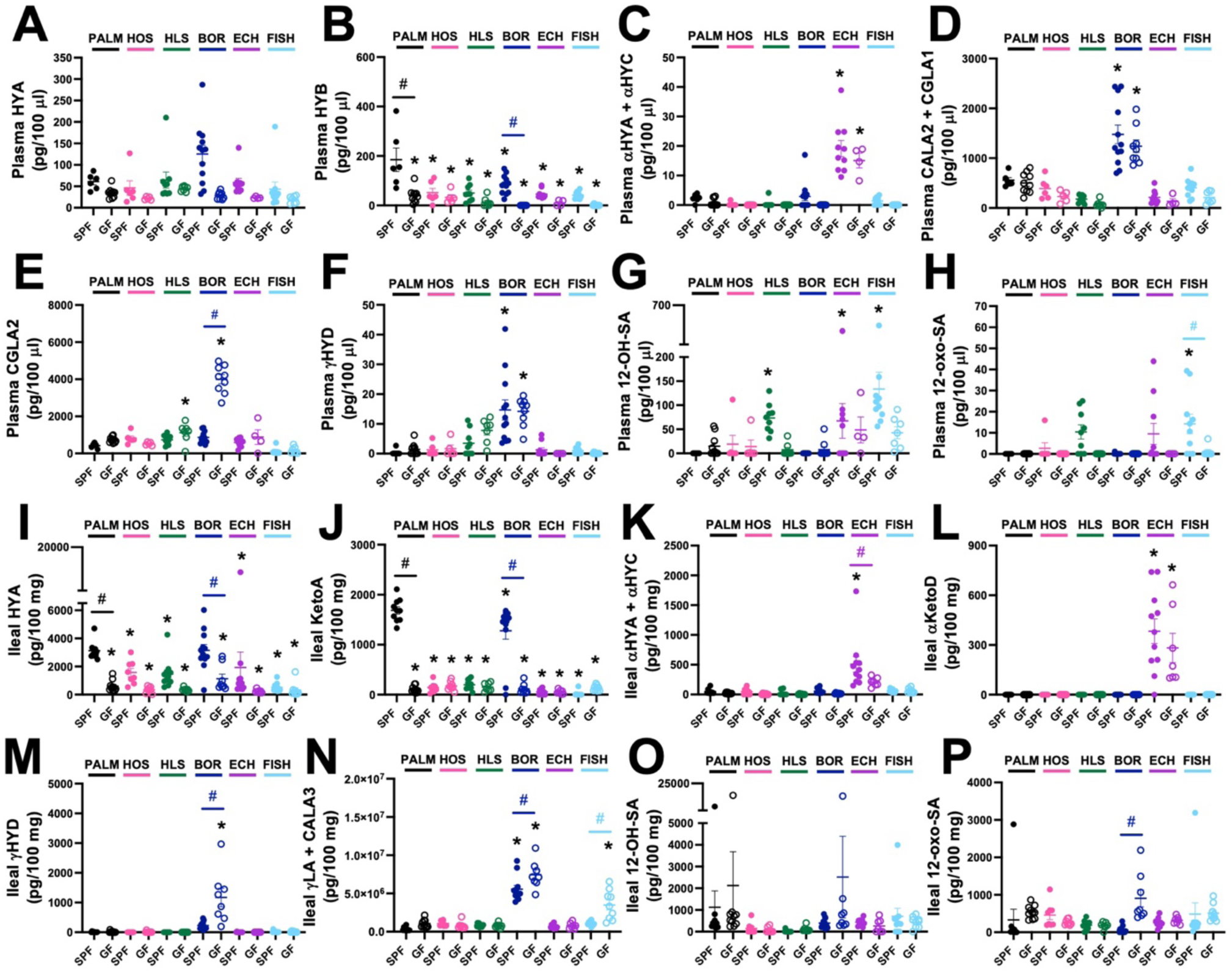
Gut Microbe-Derived Polyunsaturated Fatty Acid (PUFA) Metabolites are Altered in a Dietary Substrate-Specific Manner (*Related to* Figure 4). Starting at 6 weeks of age, either conventionally-raised specific pathogen-free (SPF) or germ-free (GF) cohorts of male C57BL/6N mice were maintained on sterile fatty acid-defined diets containing saturated (Palm), high oleic safflower (HOS), high linoleic safflower (HLS), borage (BOR), echium (ECH), or fish oils for 18 weeks. Thereafter, either plasma (panels **A-H**) or ileum tissue (panels **I-P**) were extracted to quantify bacterially-derived lipid species originating from polyunsaturated fatty acid (PUFA) metabolism. **(A-H)** Plasma levels of select bacterially-derived lipid mediators known to originate from gut microbiota-driven PUFA metabolism including **(A)** 10-hydroxy-cis12-octadecenoic acid (HYA), **(B)** 10-hydroxyoctadecanoic acid (HYB), **(C)** the sum of 10-hydroxy-cis12, cis15-octadecadienoic acid (αHYA) + 10-hydroxy-trans11, cis15-octadecadienoic acid (αHYC), **(D)** The sum of *trans*-9,*trans*-11,*cis*-15-octadecatrienoic acid (CALA2) + cis-6,cis-9,trans-11-octadecatrienoic acid (CGLA1), **(E)** cis-6,trans-9,trans-11-octadecatrienoic acid (CGLA2), **(F)** 13-hydroxy-cis6, cis9-octadecadienoic acid (γHYD), **(G)** 12-hydroxystearic acid (12-OH-SA), and **(H)** 12-oxo-stearic acid (12-oxo-SA). **(I-P)** Ileal levels of select bacterially-derived lipid mediators known to originate from gut microbiota-driven PUFA metabolism including **(I)** 10-hydroxy-cis12-octadecenoic acid (HYA), **(J)** 10-oxo-cis12-octadecenoic acid (KetoA), **(K)** the sum of 10-hydroxy-cis12, cis15-octadecadienoic acid (αHYA) + 10-hydroxy-trans11, cis15-octadecadienoic acid (αHYC), **(L)** 13-oxo-cis9, cis15-octadecadienoic acid (αKetoD), **(M)** 13-hydroxy-cis6, cis9-octadecadienoic acid (γHYD), **(N)** The sum of γ-linolenic acid (γLA) and *trans*-10,*trans*-12,*cis*-15-octadecatrienoic (CALA3), **(O)** 12-hydroxystearic acid (12-OH-SA), and **(P)** 12-oxo-stearic acid (12-oxo-SA). Data represent the mean ± S.E.M. from n=6-12 per group, and statistically significant differences (*p*<0.05) were detected using ANOVA with post-hoc Tukey-Kramer HSD for all pairs comparisons. * = significantly different than the SPF palm oil-fed control group; # = significantly different when comparing SPF and GF groups within each dietary condition.

**Figure S5.**
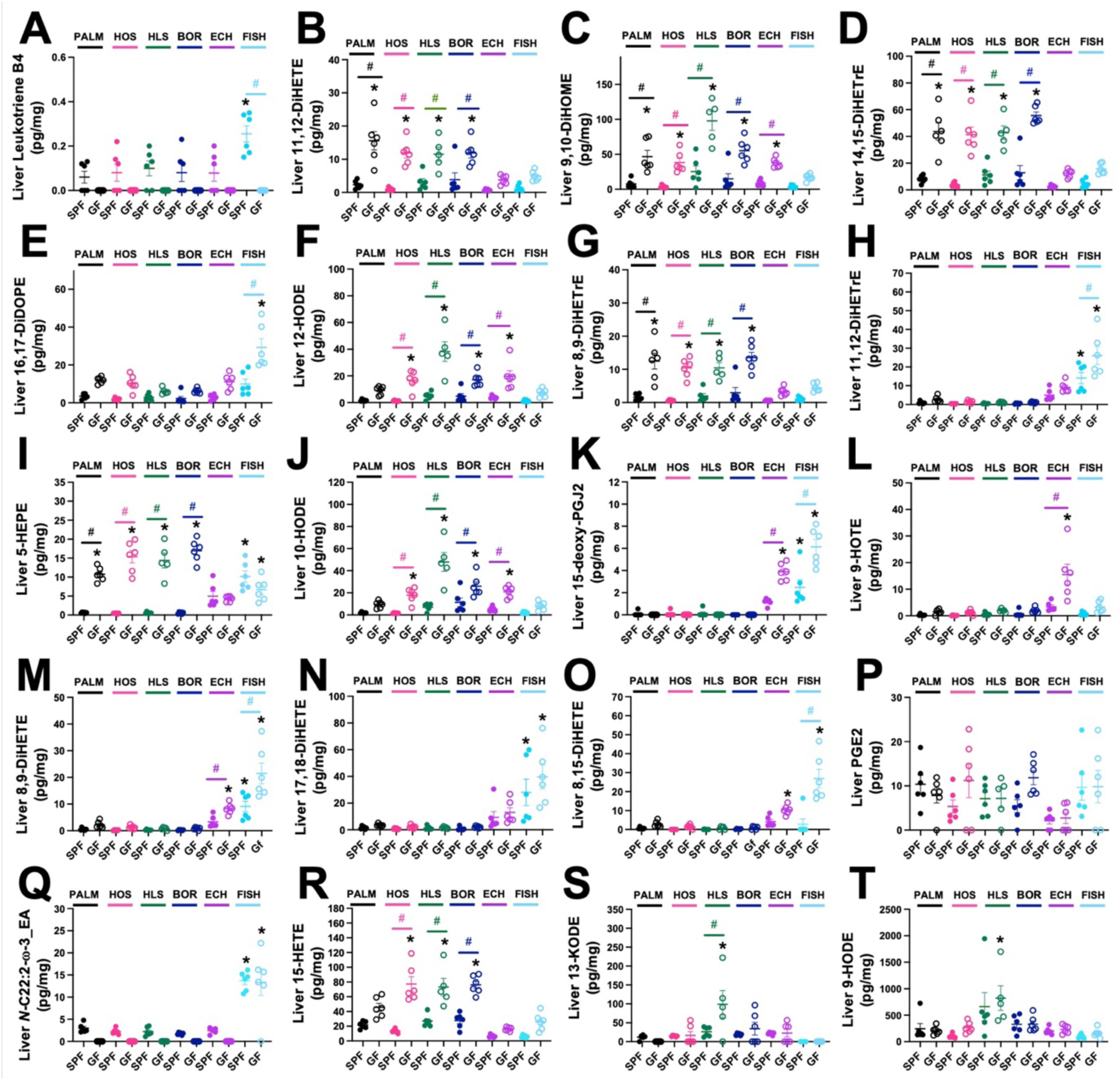
Hepatic Oxylipin Levels are Powerfully Shaped by Diet-Microbe-Host Interactions in the Mouse Metaorganism (*Related to* Figure 4). Starting at 6 weeks of age, either conventionally-raised specific pathogen-free (SPF) or germ-free (GF) cohorts of male C57BL/6N mice were maintained on sterile fatty acid-defined diets containing saturated (Palm), high oleic safflower (HOS), high linoleic safflower (HLS), borage (BOR), echium (ECH), or fish oils for 18 weeks. Thereafter, liver tissue was extracted to quantify many diverse molecular species of oxylipin-related lipids known to originate from polyunsaturated fatty acid (PUFA) substrates including: **(A)** leukotriene B4 (LTB4), **(B)** 11,12-dihydroxyicosatetraenoic acid (11,12-DiHETE), **(C)** 9,10-dihydroxy-12Z-octadecenoic acid (9,10-DiHOME), **(D) 14,15-dihydroxyeicosatrienoic acid** (14,15-DiHETrE), **(E) 16,17-dihydroxy-docosapentaenoic acid** (16,17-DiDOPE), **(F) 12-hydroxyoctadecadienoic acid (**12-HODE), **(G)** 8,9-Dihydroxy-eicosatrienoic acid. (8,9-DiHETrE), **(H)** 11,12-dihydroxyeicosatrienoic acid (11,12-DiHETrE), **(I)** 5-Hydroxyeicosapentaenoic acid (5-HEPE), **(J)** 10-Hydroxyoctadecadienoic acid (10-HODE), **(K)** 15-deoxy-prostaglandin J2 (15-deoxy-PGJ2), **(L) 9-hydroxyoctadecatrienoic acid (**9-HOTE), **(M)** 8,9-dihydroxy-eicosatetraenoic acid (8,9-DiHETE), **(N)** 17,18-dihydroxy-eicosatetraenoic acid (17,18-diHETE), **(O)** 8,15-dihydroxyeicosatetraenoic acid (8,15-DiHETE), **(P)** prostaglandin E2 (PGE2), (Q) *N*-docosadienoic acid-ethanolamide (*N*-C22:2-ω-3_EA), **(R)** 15-Hydroxyeicosatetraenoic acid (15-HETE), **(S) 13-oxooctadeca-9,11-dienoic acid** (13-KODE), and **(T)** 9-hydroxy-octadecadienoic acid (9-HODE). Data represent the mean ± S.E.M. from n=6 per group, and statistically significant differences (*p*<0.05) were detected using ANOVA with post-hoc Tukey-Kramer HSD for all pairs comparisons. * = significantly different than the SPF palm oil-fed control group; # = significantly different when comparing SPF and GF groups within each dietary condition.

**Figure S6.**
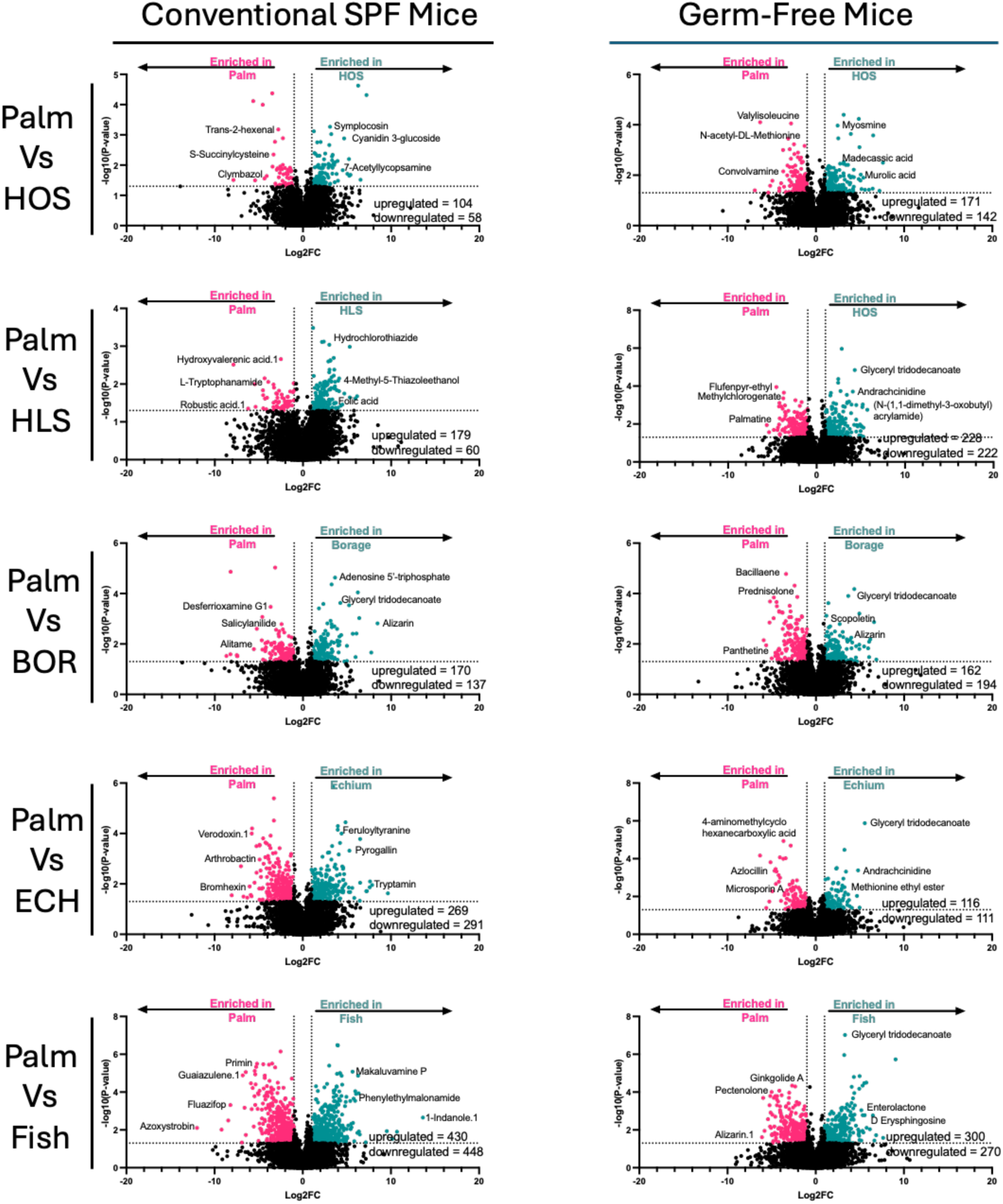
The Ability of Dietary Fatty Acids to Reorganize the Hepatic Metabolome is Altered in Germ-Free Mice (*Related to* Figures 6). Starting at 6 weeks of age, either conventionally-raised specific pathogen-free (SPF) or germ-free (GF) cohorts of male C57BL/6N mice were maintained on sterile fatty acid-defined diets containing saturated (Palm), high oleic safflower (HOS), high linoleic safflower (HLS), borage (BOR), echium (ECH), or fish oils for 18 weeks. Thereafter, liver tissue from n=6 mice per group was extracted to perform untargeted metabolomics using a hydrophilic interaction liquid chromatography (HILIC) column coupled to high resolution tandem mass spectrometry. Volcano plots are shown to compare and contrast each experimental diet group to the control palm oil-fed group either in SPF or GF mice. Significantly altered metabolites (*p*<0.05) are shown in red (decreased) or blue (increased) when compared to controls within each plots.

**Figure S7.**
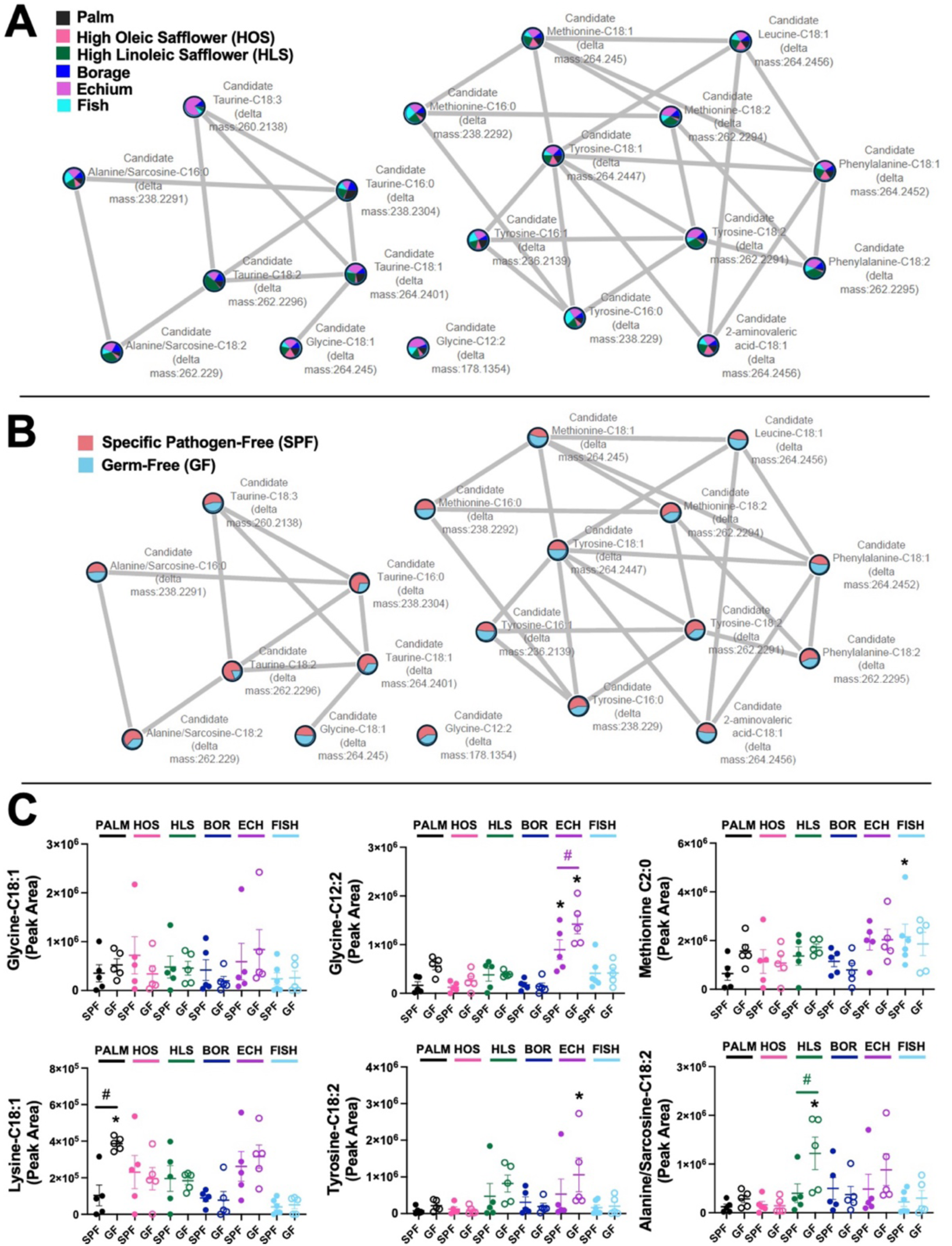
Both Dietary Fatty Acid Substrate and Resident Microbiota Impact Hepatic Levels of Diverse N-Acyl Lipids. (*Related to* Figures 6 *and Figure S6*). Starting at 6 weeks of age, either conventionally-raised specific pathogen-free (SPF) or germ-free (GF) cohorts of male C57BL/6N mice were maintained on sterile fatty acid-defined diets containing saturated (Palm), high oleic safflower (HOS), high linoleic safflower (HLS), borage (BOR), echium (ECH), or fish oils for 18 weeks. Thereafter, liver tissue from n=6 mice per group was extracted to perform untargeted metabolomics using a C18 column coupled to high resolution tandem mass spectrometry. The raw data generated were mined to identify diverse *N*-acyl lipid species. Molecular networks obtained for the N-acyl-lipids comparing across all 6 diet groups **(A)** or comparing SPF to GF cohorts **(B)**. The molecular networks were created using the feature-based molecular networking workflow within the GNPS2 environment. The nodes are annotated based on spectral similarity matches with the N-acyl lipids library. The nodes represent each MS/MS spectrum, while the edges represent their spectral similarity. The delta mass is the mass difference between the feature and Amino-containing compound. Pie charts indicate the relative abundance of ion features in each group. **(C)** The relative abundance of select N-acyl lipids show diet-microbe-host interactions.

